# Factor VIII exhibits chaperone-dependent and glucose-regulated reversible amyloid formation in the endoplasmic reticulum

**DOI:** 10.1101/2020.01.13.905190

**Authors:** Juthakorn Poothong, Anita Pottekat, Marina Siirin, Alexandre Rosa Campos, Adrienne W. Paton, James C. Paton, Jacqueline Lagunas-Acosta, Zhouji Chen, Mark Swift, Niels Volkmann, Dorit Hanein, Jing Yong, Randal J. Kaufman

**Affiliations:** Degenerative Diseases Program, SBP Medical Discovery Institute, 10901 N. Torrey Pines Rd., La Jolla CA 92037, USA; Proteomics Core Facility, SBP Medical Discovery Institute, 10901 N. Torrey Pines Rd., La Jolla CA 92037, USA; Research Centre for Infectious Diseases, Department of Molecular and Biomedical Science, University of Adelaide, Adelaide, Australia; Immunity and Pathogenesis Program, SBP Medical Discovery Institute, 10901 N. Torrey Pines Rd., La Jolla CA 92037, USA

**Keywords:** ER Stress, Clotting factor VIII, Hemophilia A, Protein misfolding, β-sheet polymerization, chaperones, ATP, BiP/GRP78, calnexin, calreticulin

## Abstract

Factor VIII (FVIII) is the coagulation factor deficient in hemophilia A, which is treated by protein replacement. Unfortunately, this regimen is costly due to the expense of producing recombinant FVIII as a consequence of its low level secretion. FVIII expression activates the endoplasmic reticulum (ER) stress response, causes oxidative stress and induces apoptosis. Importantly, little is known about the factors that cause protein misfolding and aggregation in metazoans. Here we identified intrinsic and extrinsic factors that cause FVIII to form aggregates in the ER. We show that FVIII forms amyloid-like fibrils within the ER upon increased FVIII synthesis or inhibition of glucose metabolism. Significantly, FVIII amyloids can be dissolved upon restoration of glucose metabolism to produce functional secreted FVIII. Two ER chaperones and their co-chaperones, BiP and CANX/CRT, promote FVIII solubility in the ER, where the former is also required for disaggregation. A short aggregation motif in the FVIII A1 domain (termed Aggron) is necessary and sufficient to seed β-sheet polymerization and BiP binding to this Aggron prevents amyloidogenesis. Our findings provide novel insight into mechanisms that limit FVIII secretion and ER protein folding in general and have implication for ongoing hemophilia A gene therapy clinical trials.

**Key Points:**

- FVIII forms amyloid aggregates in the ER that are dissolved in a chaperone- and glucose-dependent manner to produce secreted active FVIII.
- A short amino acid sequence in the A1 domain causes β sheet polymerization and ER chaperone BiP binding to this site prevents aggregation.

## Introduction

Hemophilia A (HA), an X chromosome-linked bleeding disorder affecting ∼10,000 males in the US, results from deficiency in coagulation FVIII, a component of the intrinsic blood clotting cascade^1,2^. Presently HA is treated by protein replacement. Although, development of recombinant-derived FVIII significantly decreased risk of adventitious viral contamination, it greatly increased the cost of treatment, partially due to the low level of FVIII secretion from recombinant mammalian host cells^3^. FVIII is synthesized and translocated into the ER lumen where only properly folded proteins traffic to the Golgi compartment. The accumulation of unfolded/misfolded proteins in the ER activates the unfolded protein response (UPR), an adaptive signaling pathway evolved to resolve ER protein misfolding^4-6^. FVIII is susceptible to misfolding in the ER and was the first native endogenous protein shown to activate the UPR by binding to the ER protein chaperone BiP/GRP78^7,8^. FVIII expression and an unresolved UPR leads to apoptosis^9^. A comprehensive understanding of the factors required for FVIII folding and secretion is unknown.

FVIII has the domain structure A1-A2-B-A3-C1-C2, where the B domain contains 18 potential N-linked glycosylation sites^1^. The A domains have amino acid homology to clotting factor V (FV) and ceruloplasmin, and structural homology to double barrel proteins, resembling the fold of cupredoxin domains^10,11^. Energy depletion, with inhibitors of mitochondrial oxidative phosphorylation and glycolysis, selectively inhibits FVIII anterograde trafficking in the early secretory pathway^12,13^, without affecting the trafficking of other proteins, including the homologous clotting factor V (FV) and von Willebrand factor (vWF), even within the same cell^12,14^. A portion of FVIII traffics to the Golgi apparatus for processing by Furin to produce a heterodimer composed of an amino-terminal ∼200kDa heavily glycosylated heavy chain (HC: A1-A2-B) in complex with a carboxy-terminal ∼80kDa light chain (LC: A3-C1-C2) linked through two copper ions in each of the A1 and A3 domains^15^ that probably stabilize the A1 and A3 domains to promote A1-A3 interaction^16,17^. Unfortunately, the molecular basis of FVIII folding and trafficking through the secretory pathway is poorly defined. Here we demonstrate that FVIII forms amyloid-like structures in the ER that can disaggregate, refold and be secreted as functional FVIII in mammalian cells. In addition, 1) a short amino acid motif, we termed Aggron, in FVIII is necessary and sufficient to seed aggregation; 2) glucose (Glc) metabolism is required to maintain FVIII solubility; and 3) two chaperone families, CANX/CRT and BiP, promote solubility, the latter is also required for FVIII disaggregation.

## Methods

### Reagents

All reagents are specified in Suppl. Information.

### Standard Methods

Plasmid construction, cell lines and treatment with metabolic inhibitors^18^ are detailed in Suppl. Information. FVIII immuno-microscopy, activity and antigen measurement, pulse-chase & IP analyses^19^, co-IPs and Western blotting, sucrose gradient sedimentation^18^, membrane filtration^20^ were standard and detailed in Suppl. Information.

#### Immunogold labeling TEM

CHO-K1 or FVIII induced H9 cells were processed for immuno-gold localization of FVIII as in Suppl. Methods.

#### Negative stain TEM

Lysates from CHO-K1 or H9 cells treated with SAHA were subjected to sucrose gradient sedimentation for FVIII IP, elution and analysis by TEM.

#### Transmission electron cryo-microscopy (cryo-EM)

IP’ed FVIII from sucrose gradient fractions of H9 cells was analyzed by cryo-EM as in Suppl. Information.

### Quantification and statistical analysis

All statistical analysis used Prism software 7. P values were calculated using one-way ANOVA. P < 0.05 was considered significant. Statistical significance in figures and legends is denoted by asterisks (^***^, p < 0.001; ^****^, p < 0.0001).

## Results

### FVIII forms amyloid-like aggregates in the ER

Newly synthesized FVIII aggregates in the cell as monitored by sucrose gradient sedimentation^18^. Immunofluorescence (IF) microscopy was used to visualize steady state FVIII aggregation two different CHO cell clones. H9 cells express a low level of human FVIII that is inducible by histone deacetylase inhibitors (HDAC) [sodium butyrate (NaB) or suberoylanilidehydroxamic acid (SAHA)]. In H9 cells, Increased FVIII expression activates the UPR and apoptosis (Fig. 1A)^8,18^. 10A1 cells express high levels of FVIII constitutively without UPR activation^19^. FVIII in H9 and 10A1 cells displayed diffuse ER localization identified by co-localization with KDEL-containing ER proteins (Fig. 1Bi; Fig. S1i). NaB induction of FVIII synthesis in H9 cells increased FVIII staining that also co-localized with the KDEL ER marker (Fig. 1Bii). Treatment of 10A1 cells or H9 cells with 2’-deoxyglucose (2DG) and sodium azide (NaN_3_) to inhibit Glc metabolism and oxidative phosphorylation, respectively, caused FVIII and KDEL proteins to co-localize to large perinuclear structures (Fig. 1Biii; Fig. S1i vs S1ii). Thioflavin-S (Thio-S), upon selective binding to β-rich structures such as amyloid, increases fluorescence intensity^21,22^. Intriguigingly, the metabolic poisons caused co-localization of Thio-S with FVIII, suggesting FVIII in the ER gained amyloid-like properties. Importantly, replacing the metabolic inhibitors with Glc-containing media for 4h, caused FVIII and KDEL proteins to resum their diffuse web-like ER co-localization and Thio-S co-staining was significantly reduced in both H9 and 10A1 cells (Fig. 1Biv; Fig. S1iii), suggesting at the histological level that FVIII aggregates disappear and traffic the secretory pathway.

**Figure 1.**
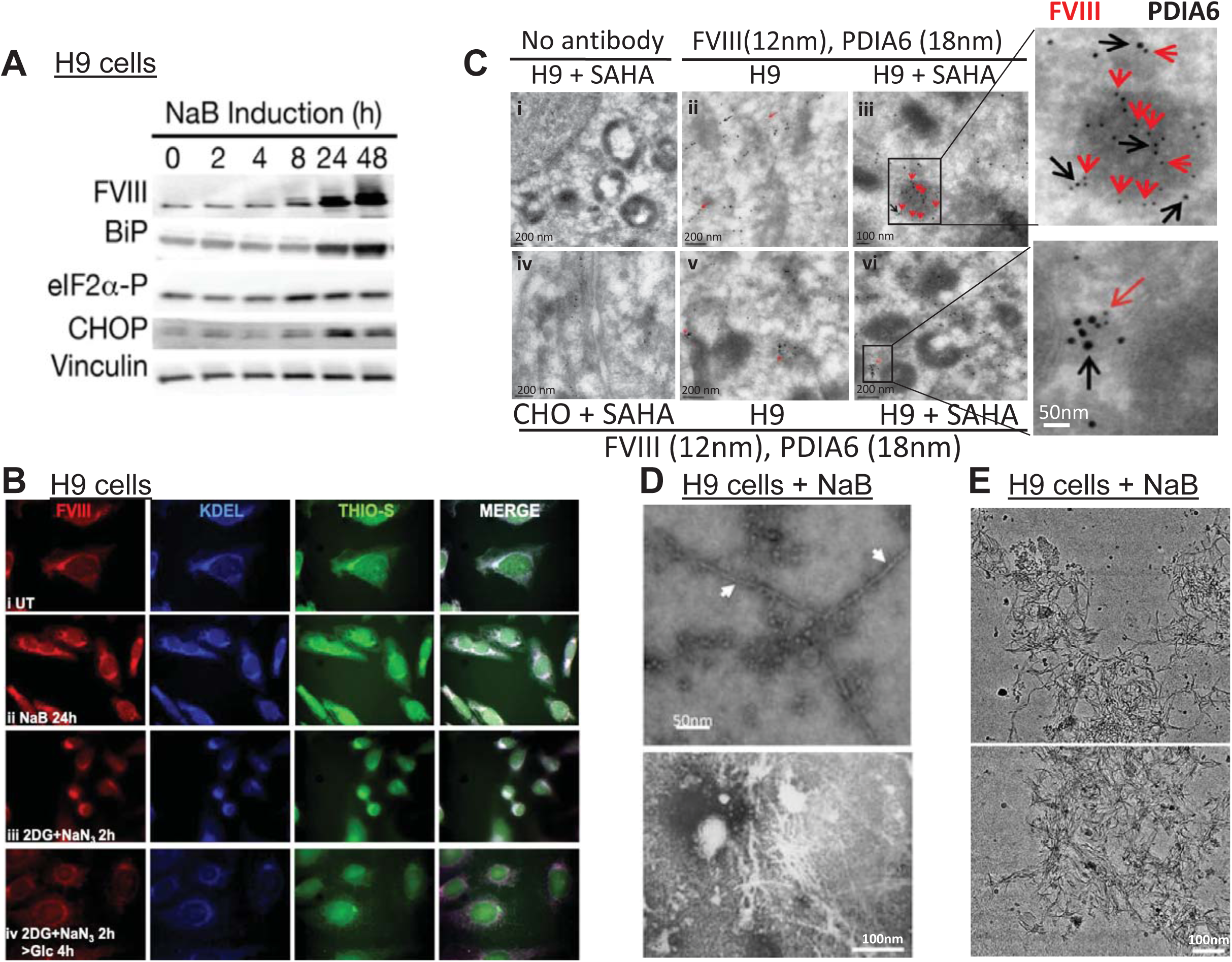
FVIII forms amyloid-like structures in the ER. **A.** Increased FVIII synthesis activates the UPR. H9 cells were treated with 5mM NaB to induce FVIII expression for increasing times and cell lysates prepared for immunoblotting with indicated antibodies. **B.** FVIII aggregates within the ER lumen. H9 cells were cultured in: **i)** Complete media; **ii)** NaB (5mM) for 24h; **iii**) Glc-free media containing 20mM 2DG and 10mM NaN_3_ for 2h or **iv**) 2DG and NaN_3_ treatment for 2h and then cells were recovered in Glc-containing media for 4h. Cells were fixed and stained with Thio-S (green) or antibodies for FVIII (red) and KDEL (blue). **C.** Co-localization of FVIII and PDIA6. CHO-K1 cells and H9 cells were treated without/with 5μM SAHA for 24h and ultra-thin sections were prepared for immunogold labeling and TEM by first staining with FVIII primary antibody using secondary antibody coupled to 12nm gold particles. Subsequently, grids were stained with PDIA6 antibody using a secondary coupled to 18nm gold particles: **i)** SAHA-treated H9 cells stained without primary antibody; **ii/v**) Untreated H9 cells; **iii/vi**) SAHA-treated H9 cells; and **iv)** SAHA-treated control CHO cells. Panels on the right show regions in **iii** and **vi** in greater detail. Red and black arrows indicate FVIII and PDIA6 particles, respectively. **D.** FVIII forms amyloid-like structures. H9 cells were treated with 5mM NaB for 24h, lysates prepared and analyzed by sucrose gradient sedimentation. FVIII in each fraction was IP’ed with FVIII-monoclonal antibody-conjugated Sepharose beads. After IP, FVIII was eluted and analyzed by negative-stain TEM. Two images are shown. Arrows indicate FVIII fibrils in the high molecular weight (HMW) sucrose fractions. **E.** Two cryo-EM images of FVIII IP’ed from the HMW fractions from H9 cells reveal dense networks composed of ∼5nm wide fibrils. Bar is 100nm.

FVIII aggregation was also analyzed by immuno-gold transmission electron microscopy (TEM). There was insignificant FVIII reactivity in controls not treated with primary antibody (Fig. 1Ci), CHO cells that do not express FVIII (Fig. 1Civ) and modest reactivity in H9 cells that were not induced to express FVIII (Fig. 1Cii,v). Upon FVIII induction in H9 cells, FVIII was detected in clusters detected by two independent FVIII antibodies coupled to 12nm gold particles (Fig. 1Ciii,vi). In addition, immunolocalization of PDIA6 by secondary antibody conjugated to 18nm gold particles demonstrated co-localization of FVIII and an ER resident protein (Fig. 1Ciii,vi and expanded on right).

The structure of purified FVIII aggregates in NaB-treated H9 cells was analyzed by conventional TEM and transmission electron cryo-microscopy (cryo-EM). After FVIII induction, cell lysates were sedimented on sucrose gradients^18^. FVIII was immunoprecipitated (IP’ed) from each fraction, eluted from the beads and analyzed by TEM and cryo-EM. Fibrils of approximately ∼5nm diameter were visible in negatively stained TEM images from the heavy molecular weight (HMW) fractions (white arrows, Fig. 1D) from NaB-treated H9 cells, likely representing aggregated FVIII. These fibrils were absent in the light fractions, presumably containing monomeric FVIII (not shown). Cryo-EM images of FVIII IP’ed from HMW fractions displayed dense networks of fibrils with diameters of 5.1±0.5nm. The fibrils appeared to interact with each other in various ways and form amyloid-like aggregates reminiscent of Poly-Q aggregates observed by cryo-EM in mammalian cells^23^ (Fig. 1E). Antibody gold labeling for FVIII further confirmed that these fibrils are composed of FVIII. Importantly, no fibrils were detected in fractions similarly prepared from CHO cells that do not express FVIII (not shown). Therefore, FVIII forms fibrils which are morphologically similar to those reported for other amyloidogenic proteins^24^.

### Aggregated FVIII disassembles and refolds into secreted functional protein

Energy depletion, with inhibitors of mitochondrial oxidative phosphorylation and glycolysis, causes FVIII aggregation and selectively inhibits FVIII trafficking to the Golgi^12,14^. Analysis of FVIII aggregation by sucrose gradient fractionation is time consuming and does not allow easy comparison between multiple variables. To examine the fate of FVIII in cells upon bioenergetic collapse and recovery we performed pulse-chase radiolabeling and IP for SDS-PAGE and autoradiography^19^, and the same samples were analyzed by a cellulose acetate (CA) filter-binding assay^20^ which retains amyloid aggregates for analysis of FVIII aggregation in a more reproducible, robust, and convenient manner. Cells were pulse-radiolabeled, treated with or without 2DG and NaN_3_ for 2h and subsequently recovered in Glc-containing medium for up to 4h for harvest of cell lysates and conditioned media. Membrane filtration and detection with FVIII antibody demonstrated immediately after treatment with 2DG and NaN_3_, which decreased cellular ATP by 5-fold (Fig. S2A,B), increased FVIII retention on CA that subsequently declined upon Glc recovery (Fig. 2Ai). FVIII antibody reactivity upon filtration through nitrocellulose (NC), which retains all cellular proteins, did not significantly change over the time course (Figs. 2A and S2A). Pulse-chase demonstrated that untreated 10A1 cells efficiently secreted FVIII at 4h after the pulse-label (Fig. 2Aii, lane 1 vs 3). In contrast, 2DG and NaN_3_ treatment prevented secretion of the pulse-labeled FVIII (Fig. 2Aii, lane 4). Upon Glc repletion, labeled FVIII appeared in the medium after 1h and accumulated to 4h (Fig. 2Aii, lanes 5-8). The total amount of labeled FVIII secreted after 4h was close to that secreted from cells that were not treated with 2DG and NaN_3_ (100% vs 87%). Notably recovery in Glc not only increased FVIII secretion (Fig. 2Aii, lane 8), but also functional FVIII activity in the medium (Fig. 2Aiii). Adding CHX during the chase recovery reduced the FVIII activity in the medium by ∼50% (Fig. 2Aiii, lane 9), indicating that ∼half of the FVIII activity was previously synthesized and was likely derived from aggregated FVIII. In addition, the decline in aggregated FVIII was not due to proteasomal or autophagic degradation (Fig. S2C). These findings support the notion that FVIII intracellular aggregates were dissolved and refolded for secretion of active FVIII.

**Figure 2.**
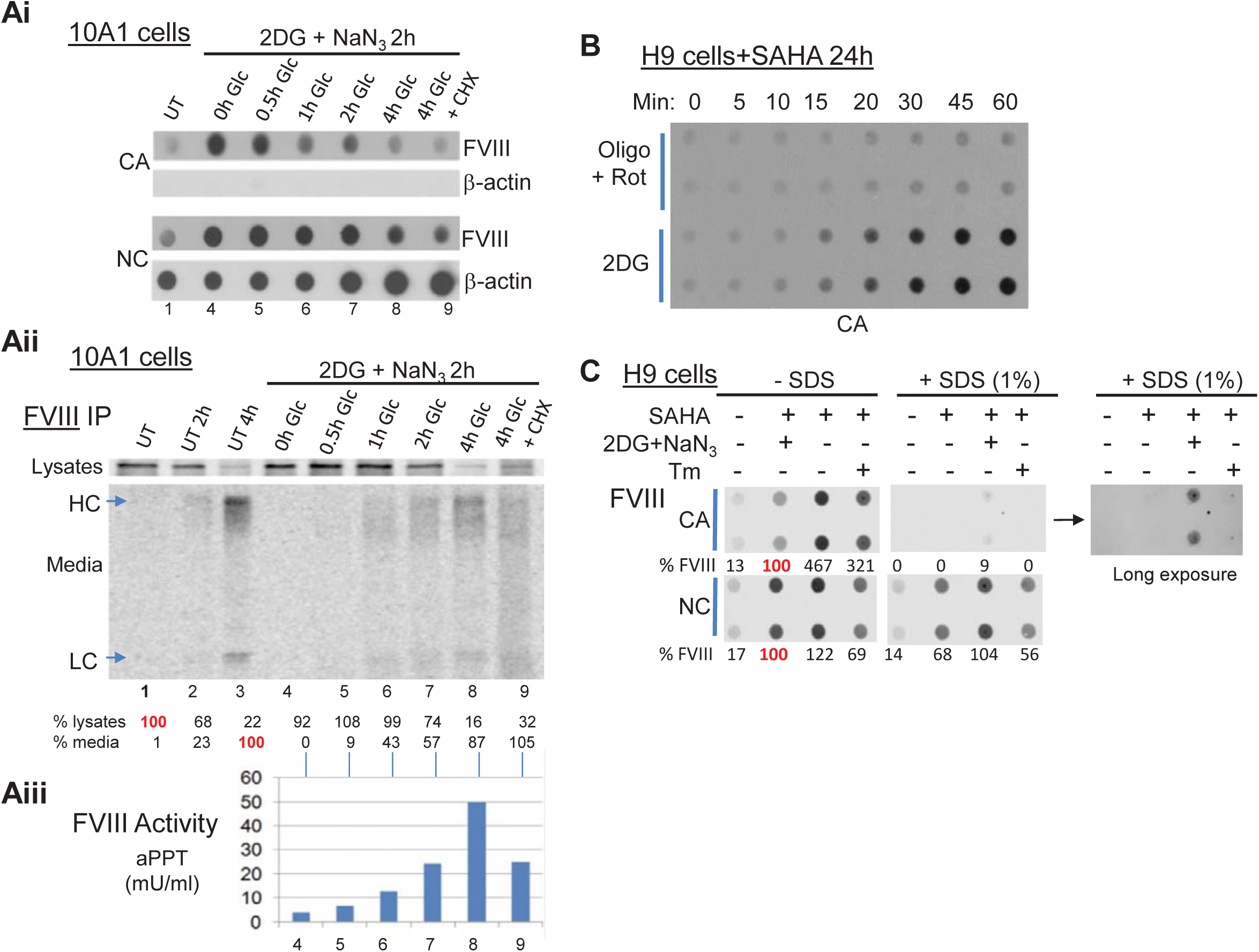
FVIII aggregation is reversible upon restoration of Glc metabolism. **A.** FVIII aggregates disassociate and refold into functional secreted FVIII. **Ai-iii.** Kinetics of FVIII aggregation and disaggregation upon altered Glc metabolism measured by membrane filtration and pulse-chase analyses. For **Ai-Aiii**, the corresponding lane numbers are indicated below. 10A1 cells were pulse-labeled for 20min with [^35^S]-Met/Cys and then chased for 20min with media containing excess unlabeled Met/Cys to compete synthesis of nascent chains (lane1). Radiolabeled cells were then treated with complete media for indicated times (**Aii.** lanes 2, 3) or Glc-free media containing 20mM 2DG and 10mM NaN_3_ for 2h (**Ai**,**ii**. lanes 4-9). Radiolabeled [^35^S]-Met/Cys cell lysates and media were harvested after 2h (lane 4) or allowed to recover in complete media (5mM Glc) from 30min to 4h (lanes 5-8), or complete media with 10μg/ml CHX (lane 9). **Ai.** At the indicated times cell lysates were analyzed by filtration on CA and NC membranes using FVIII and β-actin antibodies. **Aii.** FVIII was IP’ed from cell lysates and media for analysis by SDS-PAGE and autoradiography. FVIII heavy and light chains (HC and LC) are indicated. The percentages of radiolabeled intracellular FVIII and secreted FVIII are shown relative to intracellular FVIII in lane 1. **Aiii.** Secreted FVIII is functional by the activated Partial Thromboplastin Time (aPTT) assay. **B.** Inhibition of Glc metabolism and not oxidative phosphorylation causes FVIII aggregation. H9 cells treated with SAHA for 19h were subsequently treated with a mixture of 1μM oligomycin A (Oligo) and 1μM rotenone (Rot) or 50mM 2DG for indicated times (0-60 min). Cell lysates were filtered through CA membranes and probed with FVIII antibody. **C.** FVIII aggregation is due to hydrophobic interactions. H9 cells treated with SAHA for 22h were treated with 20mM 2DG and 10mM NaN_3_ for 2h. Treatment with Tm started at 18h after SAHA. The % FVIII aggregation on CA was normalized to FVIII on NC membranes. The longer exposure shows ∼3% of the FVIII aggregates are resistant to 1%SDS.

### Glucose metabolism promotes FVIII solubility

We then tested whether glycolysis and/or oxidative phosphorylation is required to prevent FVIII aggregation using specific inhibitors. Inhibition of Glc metabolism by 2DG immediately induced FVIII aggregation, where inhibition of oxidative phosphorylation had no effect (Fig. 2B, Fig. S2D) indicating that Glc metabolism is required to solubilize FVIII. The role of Glc metabolism in reducing FVIII aggregates is under investigation. We next tested whether FVIII aggregation involves hydrophobic interactions. The FVIII aggregates in cell lysates were solubilized with 1%SDS prior to CA filtration (Fig. 2C), although a long exposure demonstrated that ∼3% of the FVIII aggregates were resistant to 1%SDS.

### A β sheet within the FVIII A1 domain is necessary and sufficient for FVIII aggregation

Mutation of Phe309 in FVIII to Ser, the homologous residue in FV (Fig. 3A), conferred the efficient secretion of FV onto FVIII^13^. To test whether this region predisposes to aggregation, we analyzed mutants that increase (F309S; ES) or decrease [7LF-A(7 mutations of L and F to A); F306W] FVIII secretion by CA membrane retention. Expression of mutants VES and TES was reduced, thus limiting interpretation. Mutations that increased FVIII secretion (F309S and ES) reduced aggregation and those that reduced secretion increased aggregation (Fig. 3B,C), indicating this region might predispose to aggregation.

**Figure 3.**
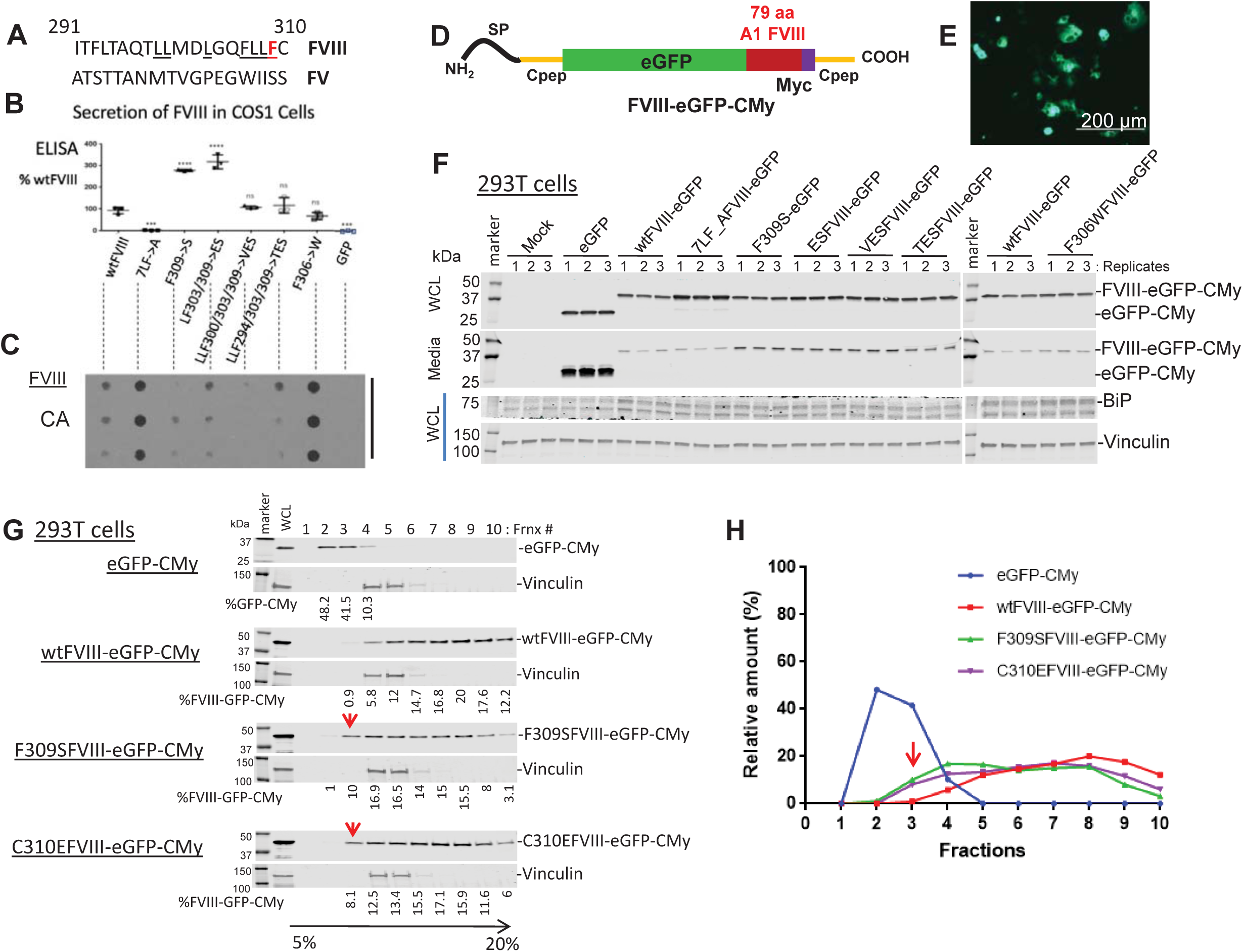
A short amino acid sequence in the A1 domain seeds FVIII aggregation. **A.** Homology is shown between aa291-310 between human FVIII and FV, with FVIII-F309 highlighted in red. **B-C.** Mutations in the FVIII A1 domain exhibit different secretion. COS-1 cells were transfected with expression vectors encoding indicated single or multiple FVIII A1 mutations. 7LF>A has 7 Phe/Leu mutations to Ala underlined in 3A. At 48h after transfection, intracellular and secreted FVIII antigen were measured by ELISA. **B.** Secreted FVIII relative to wtFVIII is shown as Mean±SD from biological triplicates. Statistical analysis was performed using one-way ANOVA: ****: p<0.0001, ***: p<0.001 compared to wtFVIII expression provided by GraphPad Prism software. **C.** Lysates from transfected cells (the same as in B) were filtered through CA membranes to measure FVIII aggregation. **D-G.** 79aa seed FVIII aggregation. **D.** Schematic structure is shown for FVIII-eGFP-CMy chimeras. The FVIII 79aa were inserted in frame downstream of eGFP and upstream of the human proinsulin C peptide (cPep) with a Myc tag (eGFP-CMy). The Myc tag is highlighted by in purple and human Cpep is in yellow. SP represents the proinsulin signal peptide. **E.** wtFVIII-eGFP-CMy transfected cells were analyzed by fluorescence microscopy. **F.** Mutations within 79aa exhibit different FVIII-eGFP-CMy secretion. Intracellular and secreted FVIII-eGFP-CMy from 293T cells expressing wtFVIII-eGFP-CMy or different mutants (wt, 7FL>A, F309S, ES, VES,TES, and F306W) was analyzed by Western blotting using Cpep antibody. The same membranes were probed with BiP and vinculin antibodies. Data shown are biological triplicates. **G-H.** wtFVIII-eGFP-CMy forms high molecular weight (HMW) complexes. Cells lysates from 293T cells expressing wtFVIII-eGFP-CMy or mutant FVIII-eGFP-CMy were subjected to 5-20% sucrose gradient sedimentation. Proteins from fractions 1-10 were analyzed by Western blot using Cpep antibody. The membrane was probed with vinculin antibody for loading control. The proportion of protein in each fraction is indicated as a percentage of total and plotted in (**H**). Red arrows indicate migration of soluble forms of F309SFVIII-eGFP-CMy and C310EFVIII-eGFP-CMy.

To test whether residues in the FVIII A1 domain are sufficient to promote aggregation, we introduced 79 residues (aa253-331) in frame into the enhanced green fluorescent protein (eGFP) fused to the human proinsulin C-peptide (Cpep) containing a cMyc tag (herein, CMy) (Fig. 3D). Antibody to Cpep conveniently recognizes all these chimeras. These 79aa were chosen because they comprise the entire copper ion binding site (Fig. S3A). These chimeras have no glycosylation sites or disulfide bonds, therefore providing the ability to disentangle FVIII aggregation from post-translational modifications. Fluorescence microscopy demonstrated significant green fluorescence in wtFVIII-eGFP-CMy expressing cells indicating insertion of these 79aa did not disrupt eGFP synthesis and folding (Fig. 3E). Western blotting of cell lysates and media demonstrated, in contrast to eGFP-CMy, insertion of the 79aa into eGFP-CMy severely inhibited secretion that was further reduced by the multiple 7LF-A mutations (Fig. 3F). Interestingly, secretion of the F309S, TES, VES and ES mutations was increased compared to wtFVIII-eGFP-CMy (Fig. 3F) indicating these 79aa are necessary and sufficient to seed aggregation. Notably expression of wtFVIII-eGFP-CMy caused ER stress monitored BiP induction in transfected cells (Fig. 3F). Thus, we propose these 79aa, termed Aggron, initiate FVIII amyloid formation.

Application of the TANGO algorithm, which predicts cross β-sheet propensity^25^, to the 79aa identified two regions that can form β-sheets (Fig. S4). Although C310S substitution did not alter the TANGO plot, F309S or C310E substitutions or the respective 79aa from FV eliminated the second β-sheet at F309. In contrast, F306W substitution, which further reduces secretion, enhanced the peak at F309. We tested the significance of these residues by expressing F309S and C310E mutations for analysis of aggregation by sucrose gradient sedimentation (Fig. 3G,H). In contrast to eGFP-CMy without FVIII sequence which migrated in light fractions 2-4, the majority of wtFVIII-eGFP-CMy migrated in heavy fractions 5-10. In contrast for both F309S and C310E, ∼10% of the FVIII-eGFP-CMy sedimented in fraction 3. Thus, these results support the validity of the TANGO analysis and show both F309 and C310 significantly contribute to FVIII aggregation.

The aggregation properties of the 79aa was further analyzed by removing eGFP to study wt and F309S FVIII-CMy in context of a smaller peptide (Fig. 4A). For controls, CMy was expressed alone or with insertions of eGFP (eGFP-CMy) or the homologous residues from FV (FV-CMy). These polypeptides were detected in transfected 293T cells by Western blotting (Fig. S3B). Cpep ELISA revealed wtFVIII-CMy was poorly secreted, where FV-CMy and F309SFVIII-CMy secretion was ∼5 to ∼10-fold greater (Fig. 4B). However, secretion of the unrelated controls CMy or GFP-CMy was significantly further increased. Inversely correlating with secretion, wtFVIII-CMy was aggregated upon CA filtration, whereas F309SFVIII-CMy aggregation was reduced to 60% (Fig. 4C). Aggregation of eGFP-CMy and FV-CMy was not detectable (Fig. 4F). Sucrose gradient sedimentation also demonstrated reduced aggregation for F309S FVIII-CMy (Fig. 4D and Fig. S5). We next tested whether the wtFVIII-CMy aggregates have properties similar to full-length wtFVIII aggregates. Sucrose gradient fractions #3 and #10 from wtFVIII-CMy expressing 293T cells were challenged with 1%SDS, 10mM DTT or β-ME prior to CA filtration. Similar to wtFVIII in H9 cells, these aggregates were sensitive to 1%SDS but resistant to DTT and β-ME (Fig. 4E, and data not shown). Together, these findings confirm that the 79aa initiate aggregation of structures that appear biochemically similar to wtFVIII.

**Figure 4.**
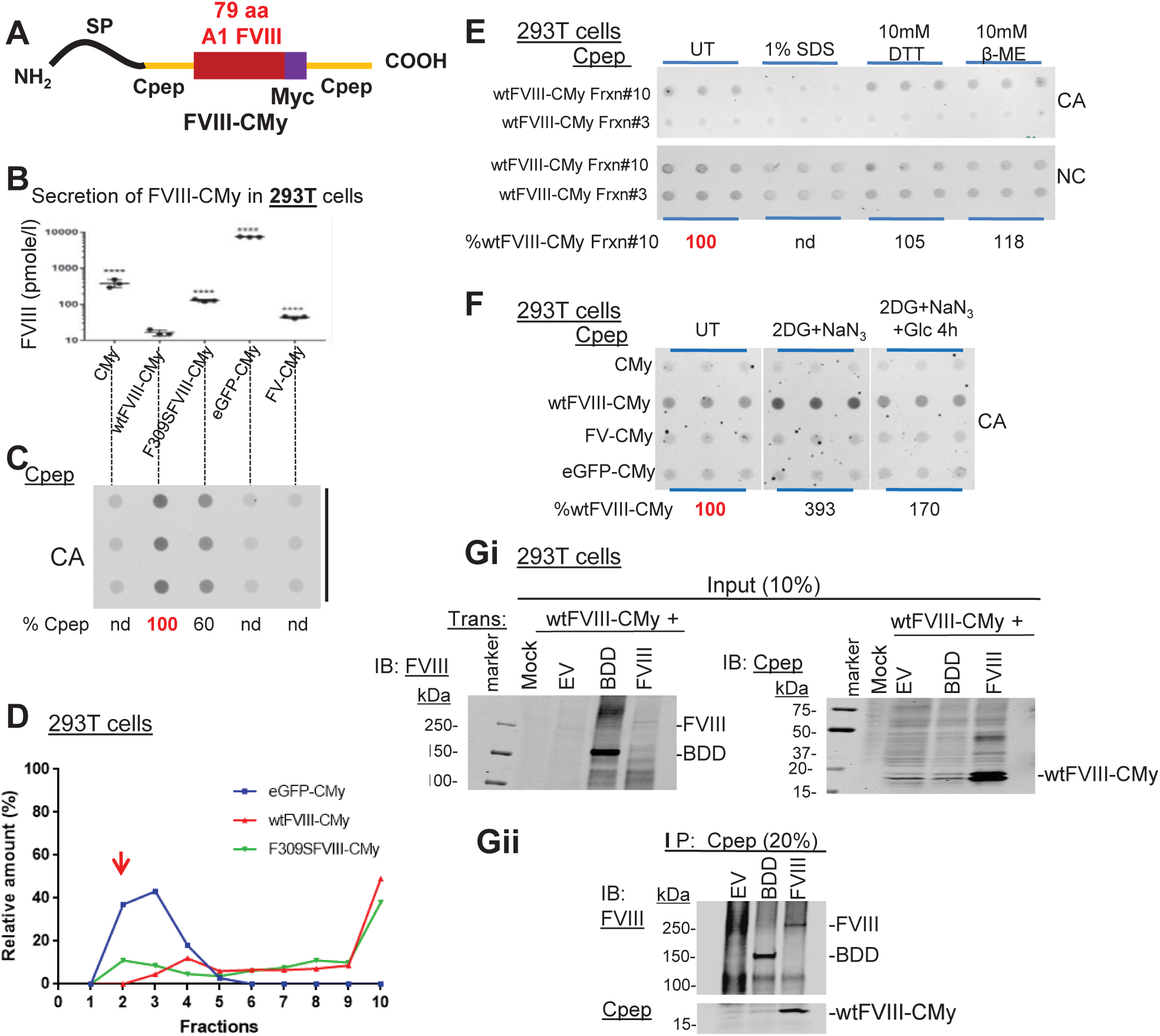
79aa from FVIII mediate C-pep reversible aggregation upon restoration of Glc metabolism. **A.** Schematic shows structure of the FVIII 79aa inserted in frame upstream of the human proinsulin C peptide (Cpep) with a Myc tag to generate wtFVIII-CMy. **B-C.** 79aa from FVIII causes Cpep aggregation. 293T cells were transfected with indicated expression plasmids: CMy, wtFVIII-CMy, F309SFVIII-CMy, GFP-CMy and FV-CMy. At 48h post-transfection, culture media and whole cell lysates (WCL) were harvested for analysis by Cpep ELISA (**B**) and aggregation by filtration onto CA membranes and probed with Cpep antibody (**C**). Data in **B** are presented as mean±SD on Log_10_ scale from biological triplicates. Statistical analysis was performed using one-way ANOVA: ****: p<0.0001 compared to wtFVIII-Cpep provided by GraphPad Prism software. **D.** wtFVIII-Cpep aggregates in HMW complexes. Cell lysates from 293T expressing wtFVIII-CMy, F309SFVIII-CMy and GFP-CMy were analyzed by 5-20% sucrose gradient sedimentation as in Fig. 3G. Protein samples from 1-10 fractions were analyzed by Western blotting using Cpep antibody. The proportion of protein in each fraction is indicated as a percentage of total. Red arrows indicate migration of soluble forms of F309SFVIII-CMy. **E.** wtFVIII-CMy aggregation is mediated by hydrophobic interactions but not disulfide cross-links. Sucrose gradient fractions 3 and 10 were treated with/without 1%SDS or with reducing agents (10mM DTT or β-mercaptoethanol (β-ME)) before CA filtration and probing with Cpep antibody. Blue bars represent technical triplicates. **F.** Aggregation of wtFVIII-CMy is reversible and requires Glc. 293T cells expressing either CMy, wtFVIII-CMy, FV-CMy or GFP-CMy were untreated (UT) or treated with Glc-free DMEM containing 20mM 2DG and 10mM NaN_3_ for 2.5h. For repletion, the 2DG and NaN_3_ containing medium was replaced with DMEM (25mM glucose) and cells were cultured 4h. Finally, cell lysates were filtered through CA and NC membranes and probed with Cpep antibody. Blue bars = technical triplicates. The % aggregated wtFVIII-CMy is relative to UT. **G.** wtFVIII-CMy specifically interacts with BDD and wtFVIII. Cell lysates from cells co-transfected with wtFVIII-CMy and wtFVIII or BDD were IP’ed with Cpep antibody. Both input (**Gi**) and Cpep IP’ed protein (**Gii**) were probed by Western using FVIII antibody (GMA) and Cpep antibody.

Since Glc addition to 2DG-treated H9 cells solubilized wtFVIII aggregates, we tested whether 2DG causes the short wtFVIII-CMy chimera to aggregate. Indeed, 2DG and NaN_3_ treatment increased wtFVIII-CMy aggregation monitored by CA filtration, with only a slight reduction in expression (Fig. 4F and Fig. S3B). These metabolic inhibitors did not significantly cause aggregation of CMy, FV-CMy or eGFP-CMy. In addition, recovery in Glc decreased wtFVIII-CMy aggregation ∼50% (Fig. 4F), consistent with observations of wtFVIII in H9 and 10A1 cells. This suggests that FVIII aggregation was not simply a consequence of altered glycosylation in response to 2DG because wtFVIII-CMy contains no N-linked glycans.

Finally, the potential of wtFVIII-CMy to form intermolecular heterodimers/oligomers with FVIII was investigated. wtFVIII-CMy and full-length wtFVIII or B domain-deleted FVIII (BDD), which aggregates similar to wtFVIII (not shown), were co-expressed in 293T cells, and their interaction was analyzed by co-IP. Surprisingly, Cpep IP of the short chimera wtFVIII-CMy efficiently pulled-down both wtFVIII and BDD (Fig. 4Gi, ii), suggesting that these 79aa interact in *trans* to alter the wtFVIII folding pathway.

### BiP and CANX/CRT prevent FVIII aggregation in the ER

Mass spectrometry of FVIII IP’ed from control CHO cells and H9 cells treated with NaB identified many ER chaperones and enzymes that selectively interact with wtFVIII (Fig. S6A,B). FVIII interactions with BiP, CANX, UGGT1, SEL1L and PDIA6 were validated by FVIII IP and Western blotting (Fig. S6C). PDIA6 also co-localized with FVIII by immuno-gold TEM (Fig. 2A).

The two most significant FVIII interactors were BiP/HSPA5 and its co-chaperones (ERdj5, ERdj3, and GRP170) and CANX/CRT with its associated factors UGGT1 and ERP57/PDIA3, raising question whether these chaperones influence FVIII aggregation. Complete inhibition of the CANX/CRT cycle with castanospermine (CST), an inhibitor of α-glucosidase (GS1 and GS2) activity, increased FVIII aggregation without altering FVIII expression (Fig. S7A-C), suggesting that CANX/CRT prevents wtFVIII aggregation, possibly by binding soluble mono-glucosylated N-linked glycans on FVIII. However, CST treatment of cells expressing the wtFVIII-CMy short chimera did not significantly increase wtFVIII-CMy aggregation (Fig. S7D,E), confirming that aggregation of the 79aa is not due to altered glycosylation because this chimera has no N-glycans.

To study the role of BiP in FVIII aggregation, we analyzed whether BiP binds to LMW and/or HMW wtFVIII species. Cell lysates from NaB-treated H9 cells were analyzed before (Fig. 5Ai) and after sucrose gradient sedimentation (Fig. 5Aii) by FVIII IP and Western blotting. Although similar amounts of wtFVIII migrated in LMW fractions 5 (L) from both untreated and NaB-treated H9 cells (Fig. 5Aii, lanes 1,3), there was a vast amount of FVIII in the HMW fraction 10 (H) (Fig. 5Aii, lane 2), with even more upon NaB induction (Fig. 5Aii, lane 4). FVIII IPs analyzed by Western blotting demonstrated detectable BiP interaction only with the LMW fraction of FVIII from NaB-treated, and not in untreated H9 cells (Fig. 5Aii, lanes1,3). BiP was not detected in the HMW fraction (Fig. 5ii, lanes 2,4), nor did we detect BiP in the post-IP HMW sucrose fraction supernatant (Fig. 5Aii, lanes 6, 8). Significantly, the majority of BiP was detected in the FVIII post-IP supernatant in the LMW fraction (Fig. 5Aii, lanes 5,7), suggesting that BiP preferentially associates with LMW FVIII.

**Figure 5.**
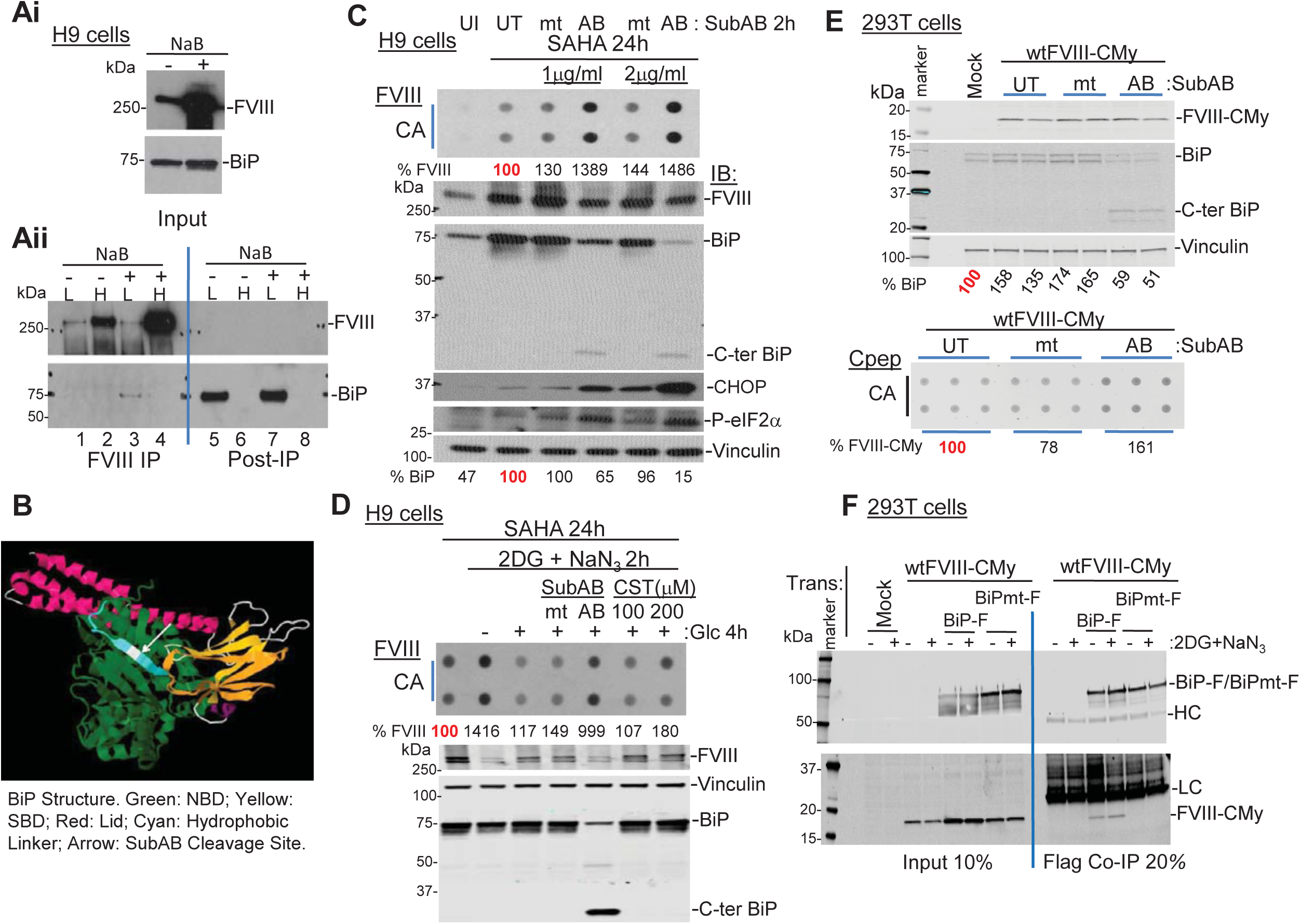
Upon Glc replenishment, BiP ATPase activity is required to solubilize wtFVIII-CMy. **A.** BiP binds LMW-FVIII. Lysates from H9 cells treated with or without NaB were subjected to sucrose gradient sedimentation and FVIII from fractions 5 (Light fraction; L) and 10 (Heavy fraction; H) were IP’ed. After elution, co-IP’ed BiP was analyzed by Western blotting. **Ai**. Western blots for FVIII and BiP are shown for cell lysates. **Aii**. Western blots are shown for FVIII IPs and post-IP supernatants. **B.** The figure shows the crystal structure of BiP in the ATP-bound conformation. The white arrow indicates the cleavage site for SubAB protease that connects the nucleotide binding domain (NBD) (Green) with the substrate binding domain (SBD) (Yellow). The C-terminal lid is indicated in pink. The model is based on PDB-6ASY, DOI: 10.2210/pdb6ASY/pdb. **C.** BiP cleavage induces wtFVIII aggregation. H9 cells were treated with 5µM SAHA for 24h and BiP cleavage was initiated by treating cells with 1 or 2μg/ml of SubAB (AB) or mutant SubA_272_B (mt) for the last 2h of SAHA treatment. **C**(top); CA membrane filtration of cell lysates shows relative levels of FVIII aggregation compared to cells not treated with SubAB (UT). **C**(bottom); Western blot is shown for cell lysates using indicated antibodies. The abundance of intact BiP (∼75kDa) was quantified relative to vinculin (loading control) and is represented as a percentage of UT. **D.** BiP is required for disassociation of FVIII aggregates. H9 cells treated with SAHA for 18h and then with 2DG and NaN_3_ for 2h in absence of Glc. Then, Glc was repleted in the presence of either 2μg/ml SubAB (AB) or SubA_272_B (mt) or CST (100 and 200μM) for 4h. Cell lysates were filtered through CA and analyzed by Western blotting. The % FVIII aggregation was quantified the FVIII signals on CA relative to Western compared to UT control. **E.** BiP inactivation induces wtFVIII-CMy chimera aggregation. 293T cells expressing wtFVIII-CMy were treated with or without SubAB or mtSubAB (2μg/ml) for 2.5h and then harvested for Western blotting and CA filtration. The relative wtFVIII-CMy Western intensities were normalized to vinculin. CA intensities were normalized to FVIII Western intensities. Black bars = biological duplicates; Blue bars = technical triplicates. **F.** wtFVIII-CMy interacts with BiP. Cells lysates from 293T cells expressing wtFVIII-CMy and BiP-Flag (BiP-F) or BiP-Flag (V461F) (BiPmt-F), a peptide-binding defective mutant, were used for Flag IP. After IP and blotting, BiP was detected with Flag antibody (top) and FVIII-CMy with cPEP antibody (bottom) to compare untreated and treated cells with 2DG and NaN_3_ for 2.5h. LC and HC represent immunoglobulin light and heavy chains of Flag antibody.

To further analyze how BiP impacts FVIII aggregation, we used the Shiga-toxic *E. coli* virulence factor SubAB that specifically and acutely cleaves at a single site in the hydrophobic linker that connects the BiP N-terminal ATPase domain with the substrate binding domain (Fig. 5B)^26,27^. As control, cells were treated with a protease mutant (SubA_272_B_mt_). Treatment of cells with 1 or 2 µg/ml of the active protease acutely cleaved ∼35% and ∼85% of BiP, respectively, and increased FVIII aggregation detected by retention on CA membranes (Fig. 5C), suggesting that intact BiP, but not cleaved BiP, prevents FVIII aggregation. BiP cleavage was associated with UPR activation by induction of P-eIF2α and CHOP (Fig. 5C), consistent with previous findings^28^.

Although the above findings indicate that BiP and CANX/CRT prevent FVIII aggregation, they do not address their role in disaggregation. Hence, we treated H9 cells with NaB for 18h with the last 2h in the presence of 2DG and NaN_3_ to induce maximal FVIII aggregation. Then, fresh media with Glc was added for 4h in the presence or absence of SubAB protease or CST. Although 2DG and NaN_3_ reduced FVIII expression, BiP cleavage prevented the Glc-dependent dissociation of FVIII aggregates (Fig. 5D). In contrast, inactivation of the CANX/CRT cycle by CST did not significantly prevent disaggregation. Collectively, we conclude that BiP, and not CANX/CRT, is required to disaggregate polymerized FVIII.

To evaluate the role of BiP in preventing FVIII aggregation, BDD-FVIII, which exhibits similar aggregation properties as wtFVIII, was co-expressed with Flag-tagged BiP (BiP-F), mutant V461FBiP-Flag (BiPmt-F), which is peptide-binding defective^29^, or Myc-tagged BiP (BiP-M)^29^. Forced expression of BiP-F or BiP-M did not alter BDD-FVIII expression, although they significantly reduced BDD-FVIII aggregation (Fig. S8). In addition, the V461F mutant BiP was partially defective in preventing BDD-FVIII aggregation. Thus, we conclude that BiP requires its peptide binding site to efficiently promote FVIII solubility in the ER.

We next tested whether BiP cleavage by SubAB impacts aggregation of the wtFVIII-CMy chimera. 293T cells expressing wtFVIII-CMy were treated with mt or wtSubAB. At 2.5h after SubAB protease addition, ∼40-50% BiP was cleaved without affecting wtFVIII-CMy expression (Fig. 5E, top). Compared to mtSubAB treatment control, wtFVIII-CMy aggregation increased ∼60% upon BiP cleavage (Fig. 5E, bottom). Although SubAB inhibits secretion of immunoglobulin^30^, secretion of eGFP-CMy was not reduced (Fig. S9A,B), indicating the secretory pathway remained intact. Collectively, these findings show that intact BiP prevents aggregation of wtFVIII-CMy through interaction with the BiP peptide binding site.

### BiP interacts with wtFVIII-CMy to prevent aggregation

We tested whether binds to the 79aa by co-expression of wtFVIII-CMy with BiP-F or the peptide-binding defective BiPmt-F^29^. Indeed, Flag IP of BiP co-IP’ed wtFVIII-CMy (Fig. 5F), suggesting a direct interaction between BiP and the 79aa of FVIII. In contrast, wtFVIII-CMy did not co-IP with mutant BiPmt-F. However, treatment with 2DG and NaN_3_ did not significantly increase the BiP-wtFVIII-CMy interaction, consistent with the notion that SubAB toxin cleavage of BiP induces wtFVIII-CMy aggregation, as we could not detect BiP binding to HMW FVIII aggregates (Fig. 5A). However, mutations that reduce FVIII-CMy aggregation did not detectably alter BiP interaction measured by co-IP (not shown). It is likely that BiP binds multiple sequences so mutations that reduce aggregation do not alter BiP interaction, as previously suggested^31,32^. These findings support the conclusion that BiP binds 79aa in wtFVIII-CMy to prevent aggregation.

## Discussion

### FVIII forms reversible amyloid-like aggregates in the ER

FVIII is poorly secreted due to retention in the ER. Here we provide significant insight into the mechanism for FVIII retention in the ER. We demonstrated that FVIII forms amyloid-like structures in the ER when expressed constitutively and to a greater extent upon increased synthesis. In addition, inhibition of Glc metabolism, but not oxidative phosphorylation, causes FVIII aggregation and reduces secretion. These aggregates were characterized by: i) sucrose gradient fractionation of cell lysates, FVIII pulse-chase labeling and IP, negative stain TEM and cryo-EM which identified ∼5nm fibrils and fibril networks only in the very HMW fractions; ii) FVIII co-localization with Thio-S in the ER; iii) FVIII retention on CA membranes; iv) immuno-EM showing FVIII clustering with the ER-localized PDIA6; and v) sensitivity to 1%SDS, although a small fraction (∼3%) was resistant, all indicative of amyloid-like fibrils. Other than immunoglobulin light and heavy chains^33^, this represents a unique example of amyloid-like aggregates formed in the ER. Remarkably, unlike other amyloids found in the ER, FVIII aggregates were reversible upon Glc supplementation and were recovered as functional FVIII in the medium. To our knowledge this is the first example of reversible amyloid-like aggregation and proper folding and secretion of a protein in the ER of metazoans.

### BiP and CANX/CRT prevent FVIII aggregation

Mass Spectrometry demonstrated FVIII interaction with BiP and CANX/CRT. Here, we show CANX/CRT interaction reduces wtFVIII aggregation, similar to other glycoproteins^34,35^. BiP preferentially binds to LMW FVIII and not HMW FVIII aggregates. BiP inactivation increased FVIII aggregation and forced BiP expression reduced FVIII aggregation. Thus, BiP is required to prevent FVIII aggregation and/or to mediate disaggregation. It was shown that increased BiP expression inhibits wtFVIII secretion^14^. However, contrary to what we expected, increased BiP expression prevented wtFVIII aggregation and retained soluble LMW FVIII in the ER. From these findings, we propose a model for BiP function in preventing FVIII aggregation (Fig. 6).

**Figure 6.**
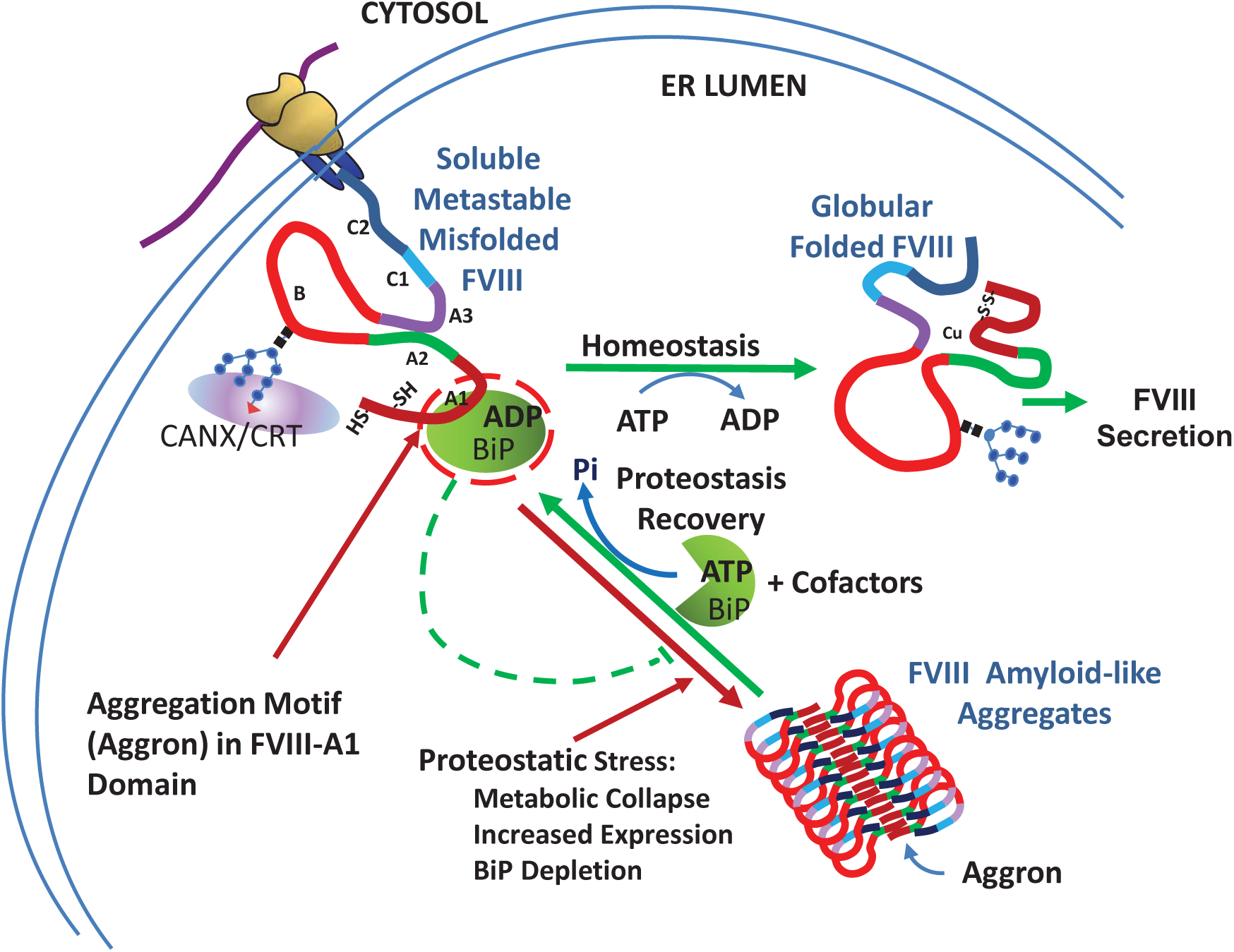
Model depicts mechanism for reversible aggregation of FVIII. Once nascent FVIII (A1-A2-B-A3-C1-C2) translocates into the ER lumen, it undergoes disulfide bond formation and N-linked core glycosylation. Glycosylated FVIII subsequently interacts with CANX/CRT and BiP, the latter at a short sequence aggregating motif (Aggron) in the A1 domain to facilitate folding by stabilizing soluble misfoflded FVIII. Upon release from CANX/CRT correctly folded FVIII (soluble) traffics to the Golgi compartment. Upon Glc supplementation, ATP hydrolysis by BiP stimulates FVIII disaggregation and release for folding and secretion as a functional clotting factor. Under proteostatic stress, such as increased FVIII expression, metabolic collapse or BiP depletion, FVIII misfolds and forms amyloid-like fibrils due to anti-parallel β-sheet polymerization initiated by the FVIII A1 domain. The Aggron and BiP binding site in the A1 domain overlap, indicated by the red circle. Red lines depict pathways of aggregation and green lines depict productive folding.

### Identification of the β-aggregate seeding region in the FVIII A1 domain

To further characterize FVIII aggregation, we identified 79aa in the A1 domain that seed β-aggregation. Expression of a chimeric polypeptide (wtFVIII-eGFP-CMy or wtFVIII-CMy) recapitulated the major aspects of wtFVIII aggregation including: i. wtFVIII-CMy aggregated in response to glycolysis inhibition; and ii. wtFVIII-CMy aggregated in response to BiP cleavage by SubAB cytotoxin. In addition, we show that the 79aa motif interacts in *trans* with wtFVIII and BDD-FVIII as well as with the peptide binding site on BiP. We propose BiP binding to the FVIII aggregate seeding region, Aggron, prevents aggregation, consistent with previous reports of other proteins^31,36,37^. As BiP prevents FVIII aggregation, it is interesting to note that three BiP co-chaperones ERdj3, ERdj5, and GRP170, the latter two which interact with aggregation prone sequences^31^, were also identified as significant interactors with wtFVIII. Although previous studies linked co-chaperone interactions with ER-associated degradation (ERAD), FVIII is not significantly degraded by ERAD, but rather is secreted. Present studies are testing the significance of these co-chaperone interactions. Importantly, without the complication from glycosylation sites and disulfide bonds, wtFVIII-CMy did not aggregate upon inhibition of the CANX/CRT cycle, implicating the glycans important for solubility. Finally, these observations of FVIII aggregation were cell-type independent as they were observed in CHO, 293T, COS-1 as well as hepatocytes *in vivo* (not shown).

### Can FVIII aggregation impact human HA gene therapy?

At least three HA gene therapy clinical trials are in progress delivering BDD-FVIII in adeno-associated viral vectors to hepatocytes (https://clinicaltrials.gov/ct2/results?term=hemophilia+a+gene+therapy). One report demonstrated significant success in 9 men with severe HA at 1 year after virus delivery to obtain 100% FVIII levels in the blood^38^. However, a more recent update indicated the FVIII levels in the high dose cohort (6e^13^ vector genomes/kg) decreased ∼50% after year 2^39^, which is unusual compared to the stable expression observed in other hepatocyte-directed AAV gene therapy trials^40,41^. All the properties of wtFVIII misfolding and aggregation also exist for the BDD-FVIII that is used in the present gene therapy clinical studies^9^. It should be considered that the decline in FVIII expression may reflect toxicity of aggregated BDD-FVIII in hepatocytes. In addition, endogenous FVIII is not likely expressed in hepatocytes, but rather in Kupffer cells and sinusoidal endothelial cells^42-44^ which may harbor machinery to efficiently fold FVIII. We also observed aggregation of BDD-FVIII in hepatocytes of mice that received BDD-FVIII DNA expression vectors, that was not observed for F309SFVIII (^9^and data not shown). Recently, it was shown that ER protein misfolding in hepatocytes with subsequent inflammation caused by a high fat diet in mice can initiate non-alcoholic fatty liver disease (NAFLD) with progression to non-alcoholic steatohepatitis (NASH) and hepatocellular carcinoma^45^. Thus, it will be important to monitor liver function in the HA gene therapy studies.

Importantly, why does FVIII harbor an Aggron in the A1 domain? Intriguingly, the Aggron includes Cys310 which ligands Cu^+^ in functional wtFVIII^15^. Of all single amino acid variants that reduce FVIII aggregation and improve secretion, F309S was the most effective. Previously, it was demonstrated that F309SFVIII exhibits clotting factor specific activity, thermal denaturation and thrombin activation indistinguishable from wtFVIII^13^. However, F309S FVIII was 10-fold more sensitive to EDTA inactivation, which presumably extracts Cu^+^. Thus, we propose this Aggron produces an anti-parallel β-sheet hairpin between the 7^th^ and 8^th^ β sheets in the double β barrel that stabilizes the C310-Cu^+^ interaction. Because F309SFVIII and F309S BDD-FVIII are functionally identical to wtFVIII and BDD-FVIII, respectively, do not aggregate or activate the UPR and are secreted efficiently, they should be considered a safe alternative to wtFVIII or BDD-FVIII for HA gene therapy.

## Supporting information

Poothong et al 2020

## Acknowledgements

We thank Dr. David Ron for kindly providing BiP expression vectors. Dr. Peter Arvan and Ming Liu for providing the pTargetT-hProCpepMyc and pTarget-hProCpepeGFP vector. Baxter Corp. (Deerfield, IL) kindly provided FVIII antibody-conjugated sepharose beads. We gratefully thank: Dr. Michael Cunningham for performing the TEM of FVIII fibrils (Fig. 1D); Dr. Lisa Giles for immunofluorescence analyses and establishing the CA filter-binding assay (Fig. 1B and S2); and Dr. Jyoti Malhotra for performing pulse-chase radiolabeling and IP (Fig. 2Aii). Immunogold labeling and TEM was performed at the UCSD Cellular & Molecular Medicine Electron Microscopy Facility. We thank the Univ. Michigan transmission electron microscopy lab for negative stained purified FVIII fibrils. We thank Dr. Peter Arvan for critical review of this manuscript. We acknowledge support from the NIH/NCI grants R01HL052173, R37DK042394, R01DK113171, R01DK103185, R24DK110973, R01CA198103, R01AG062190 and the SBP NCI Cancer Center Grant P30 CA030199. NIH Grants R01GM119948 (DH, NV) and S10-OD012372(DH) supported the cryo-EM studies. RJK is a member of the UCSD Diabetes Research Center (P30 DK063491) and an Adjunct Professor in the Department of Pharmacology, UCSD.

## Authorship Contributions

JP performed all studies on FVIII-CMy and FVIII-eGFP-CMy, experimental design and manuscript preparation.

ZC for expert consultation and manuscript preparation.

JY for sucrose gradient analyses, BiP interaction, useful discussions, and manuscript preparation.

ARC performed Mass Spec analyses and bioinformatics.

AP performed Mass Spec analyses and experiments to validate significance of FVIII interacting proteins.

JY, MS, DH and NV performed the cryo-EM sample preparation and analysis. DH and NV provided the financial support for the cryo-EM studies.

JLA provided technical support.

MS performed studies involving CA and NC membrane filtration, analysis of FVIII secretion and intracellular ATP level.

AWP and JCP provided essential reagents.

RJK directed the studies, assured validity of findings, wrote the manuscript and provided financial support for the studies.

## References

1. Toole JJ, Knopf JL, Wozney JM, Sultzman LA, Buecker JL, Pittman DD, Kaufman RJ, Brown E, Shoemaker C, Orr EC, et al. Molecular cloning of a cdna encoding human antihaemophilic factor. Nature. 1984;312:342–347.

2. Vehar GA, Keyt B, Eaton D, Rodriguez H, O’Brien DP, Rotblat F, Oppermann H, Keck R, Wood WI, Harkins RN, et al. Structure of human factor viii. Nature. 1984;312:337–342.

3. Kaufman RJ. Genetic engineering of factor viii. Nature. 1989;342:207–208

4. Walter P, Ron D. The unfolded protein response: From stress pathway to homeostatic regulation. Science. 2011;334:1081–1086.

5. Mori K. The unfolded protein response: The dawn of a new field. Proc Jpn Acad Ser B Phys Biol Sci. 2015;91:469–480.

6. Wang M, Kaufman RJ. Protein misfolding in the endoplasmic reticulum as a conduit to human disease. Nature. 2016;529:326–335.

7. Dorner AJ, Bole DG, Kaufman RJ. The relationship of n-linked glycosylation and heavy chain-binding protein association with the secretion of glycoproteins. J Cell Biol. 1987;105:2665–2674.

8. Dorner AJ, Wasley LC, Kaufman RJ. Increased synthesis of secreted proteins induces expression of glucose-regulated proteins in butyrate-treated chinese hamster ovary cells. J Biol Chem. 1989;264:20602–20607.

9. Malhotra JD, Miao H, Zhang K, Wolfson A, Pennathur S, Pipe SW, Kaufman RJ. Antioxidants reduce endoplasmic reticulum stress and improve protein secretion. Proc Natl Acad Sci U S A. 2008;105:18525–18530.

10. Zaitsev VN, Zaitseva I, Papiz M, Lindley PF. An x-ray crystallographic study of the binding sites of the azide inhibitor and organic substrates to ceruloplasmin, a multi-copper oxidase in the plasma. J Biol Inorg Chem. 1999;4:579–587.

11. Jenny RJ, Pittman DD, Toole JJ, Kriz RW, Aldape RA, Hewick RM, Kaufman RJ, Mann KG. Complete cdna and derived amino acid sequence of human factor v. Proc Natl Acad Sci U S A. 1987;84:4846–4850.

12. Dorner AJ, Wasley LC, Kaufman RJ. Protein dissociation from grp78 and secretion are blocked by depletion of cellular atp levels. Proc Natl Acad Sci U S A. 1990;87:7429–7432.

13. Swaroop M, Moussalli M, Pipe SW, Kaufman RJ. Mutagenesis of a potential immunoglobulin-binding protein-binding site enhances secretion of coagulation factor viii. J Biol Chem. 1997;272:24121–24124.

14. Dorner AJ, Wasley LC, Kaufman RJ. Overexpression of grp78 mitigates stress induction of glucose regulated proteins and blocks secretion of selective proteins in chinese hamster ovary cells. EMBO J. 1992;11:1563–1571.

15. Tagliavacca L, Moon N, Dunham WR, Kaufman RJ. Identification and functional requirement of cu(i) and its ligands within coagulation factor viii. J Biol Chem. 1997;272:27428–27434.

16. Ngo JC, Huang M, Roth DA, Furie BC, Furie B. Crystal structure of human factor viii: Implications for the formation of the factor ixa-factor viiia complex. Structure. 2008;16:597–606.

17. Shen BW, Spiegel PC, Chang CH, Huh JW, Lee JS, Kim J, Kim YH, Stoddard BL. The tertiary structure and domain organization of coagulation factor viii. Blood. 2008;111:1240–1247.

18. Tagliavacca L, Wang Q, Kaufman RJ. Atp-dependent dissociation of non-disulfide-linked aggregates of coagulation factor viii is a rate-limiting step for secretion. Biochemistry. 2000;39:1973–1981.

19. Kaufman RJ, Wasley LC, Dorner AJ. Synthesis, processing, and secretion of recombinant human factor viii expressed in mammalian cells. J Biol Chem. 1988;263:6352–6362.

20. Wanker EE, Scherzinger E, Heiser V, Sittler A, Eickhoff H, Lehrach H. Membrane filter assay for detection of amyloid-like polyglutamine-containing protein aggregates. Methods enzymol. Academic Press; 1999:375–386.

21. Wolfe LS, Calabrese MF, Nath A, Blaho DV, Miranker AD, Xiong Y. Protein-induced photophysical changes to the amyloid indicator dye thioflavin t. Proc Natl Acad Sci U S A. 2010;107:16863–16868.

22. Groenning M. Binding mode of thioflavin t and other molecular probes in the context of amyloid fibrils-current status. J Chem Biol. 2010;3:1–18.

23. Bauerlein FJB, Saha I, Mishra A, Kalemanov M, Martinez-Sanchez A, Klein R, Dudanova I, Hipp MS, Hartl FU, Baumeister W, Fernandez-Busnadiego R. In situ architecture and cellular interactions of polyq inclusions. Cell. 2017;171:179–187 e110

24. Shirahama T, Cohen AS. Structure of amyloid fibrils after negative staining and high-resolution electron microscopy. Nature. 1965;206:737–738.

25. Fernandez-Escamilla AM, Rousseau F, Schymkowitz J, Serrano L. Prediction of sequence-dependent and mutational effects on the aggregation of peptides and proteins. Nat Biotechnol. 2004;22:1302–1306.

26. Paton AW, Beddoe T, Thorpe CM, Whisstock JC, Wilce MC, Rossjohn J, Talbot UM, Paton JC. Ab5 subtilase cytotoxin inactivates the endoplasmic reticulum chaperone bip. Nature. 2006;443:548–552.

27. Yang J, Zong Y, Su J, Li H, Zhu H, Columbus L, Zhou L, Liu Q. Conformation transitions of the polypeptide-binding pocket support an active substrate release from hsp70s. Nat Commun. 2017;8:1201.

28. Wolfson JJ, May KL, Thorpe CM, Jandhyala DM, Paton JC, Paton AW. Subtilase cytotoxin activates perk, ire1 and atf6 endoplasmic reticulum stress-signalling pathways. Cell Microbiol. 2008;10:1775–1786.

29. Preissler S, Chambers JE, Crespillo-Casado A, Avezov E, Miranda E, Perez J, Hendershot LM, Harding HP, Ron D. Physiological modulation of bip activity by trans-protomer engagement of the interdomain linker. eLife. 2015;4:e08961.

30. Hu CC, Dougan SK, Winter SV, Paton AW, Paton JC, Ploegh HL. Subtilase cytotoxin cleaves newly synthesized bip and blocks antibody secretion in b lymphocytes. J Exp Med. 2009;206:2429–2440.

31. Behnke J, Mann MJ, Scruggs F-L, Feige MJ, Hendershot LM. Members of the hsp70 family recognize distinct types of sequences to execute er quality control. Mol Cell. 2016;63:739–752.

32. Knarr G, Modrow S, Todd A, Gething MJ, Buchner J. Bip-binding sequences in hiv gp160. Implications for the binding specificity of bip. J Biol Chem. 1999;274:29850–29857.

33. Brumshtein B, Esswein SR, Sawaya MR, Rosenberg G, Ly AT, Landau M, Eisenberg DS. Identification of two principal amyloid-driving segments in variable domains of ig light chains in systemic light-chain amyloidosis. J Biol Chem. 2018;293:19659–19671.

34. Ferris SP, Jaber NS, Molinari M, Arvan P, Kaufman RJ. Udp-glucose:Glycoprotein glucosyltransferase (uggt1) promotes substrate solubility in the endoplasmic reticulum. Mol Biol Cell. 2013;24:2597–2608.

35. Vassilakos A, Cohen-Doyle MF, Peterson PA, Jackson MR, Williams DB. The molecular chaperone calnexin facilitates folding and assembly of class i histocompatibility molecules. EMBO J. 1996;15:1495–1506.

36. Ushioda R, Hoseki J, Araki K, Jansen G, Thomas DY, Nagata K. Erdj5 is required as a disulfide reductase for degradation of misfolded proteins in the er. Science. 2008;321:569–572.

37. Dong M, Bridges JP, Apsley K, Xu Y, Weaver TE. Erdj4 and erdj5 are required for endoplasmic reticulum-associated protein degradation of misfolded surfactant protein c. Mol Biol Cell. 2008 Jun;19(6):2620–30. doi: 10.1091/mbc.e1007-1007-0674.

38. Rangarajan S, Walsh L, Lester W, Perry D, Madan B, Laffan M, Yu H, Vettermann C, Pierce GF, Wong WY, Pasi KJ. Aav5-factor viii gene transfer in severe hemophilia a. N Engl J Med. 2017;377:2519–2530.

39. Rangarajan S, Kim, B., Lester, W. Achievement of normal fviii activity following gene transfer with valoctocogene roxaparvovec [bmn 2017]: Long-term efficacy and safety results in patients with severe haemophilia a. WFH 2018 World Congress. 2018.

40. Niemeyer GP, Herzog RW, Mount J, Arruda VR, Tillson DM, Hathcock J, van Ginkel FW, High KA, Lothrop CD, Jr. Long-term correction of inhibitor-prone hemophilia b dogs treated with liver-directed aav2-mediated factor ix gene therapy. Blood. 2009;113:797–806.

41. Nathwani AC, Reiss U, Tuddenham E, Chowdary P, McIntosh J, Riddell A, Pie J, Mahlangu JN, Recht M, Shen Y-M, et al. Adeno-associated mediated gene transfer for hemophilia b:8 year follow up and impact of removing “empty viral particles” on safety and efficacy of gene transfer. Blood. 2018;132:491–491.

42. Kumaran V, Benten D, Follenzi A, Joseph B, Sarkar R, Gupta S. Transplantation of endothelial cells corrects the phenotype in hemophilia a mice. J Thromb Haemost. 2005;3:2022–2031.

43. Shahani T, Covens K, Lavend’homme R, Jazouli N, Sokal E, Peerlinck K, Jacquemin M. Human liver sinusoidal endothelial cells but not hepatocytes contain factor viii. J Thromb Haemost. 2014;12:36–42.

44. Everett LA, Cleuren AC, Khoriaty RN, Ginsburg D. Murine coagulation factor viii is synthesized in endothelial cells. Blood. 2014;123:3697–3705.

45. Nakagawa H, Umemura A, Taniguchi K, Font-Burgada J, Dhar D, Ogata H, Zhong Z, Valasek MA, Seki E, Hidalgo J, Koike K, Kaufman RJ, Karin M. Er stress cooperates with hypernutrition to trigger tnf-dependent spontaneous hcc development. Cancer Cell. 2014;26:331–343.

