## Supplementary material for "Factor VIII exhibits chaperone-dependent and glucose-regulated reversible amyloid formation in the endoplasmic reticulum": Poothong et al 2020

#### Supplemental Methods

##### Reagents

FVIII-deficient and normal pooled human plasma were obtained from George King Biomedical (Overland Park, KS). Activated partial thromboplastin (automated aPTT reagent) was purchased from General Diagnostics OrganonTeknika (Durham, NC). Vorinostat (SAHA) (cat#4652) and castanospermine (CST) (cat# 0759) were purchased from Fisher Scientific. FVIII:C-ELISA was purchased from Affinity Biologicals. Ultrasensitive Cpep ELISA kit (cat#10-1141-01) was purchased from Mercodia AB. 3-MA, 2DG, and $NaN_3$ were obtained from Sigma Aldrich. Antibodies to BiP (cat#C50B12), P-eIF2α (cat#3597), LC3 (cat#4599), ATF4 (cat#11815), ERP44 (cat#2886s), Myc (9B11) (cat#2276), vinculin (cat#V9131) and β actin (cat#3700s) were from Cell Signaling Technologies. Antibodies to CHOP (cat#sc-575), ERGIC53 (cat# sc-365158) and KDEL (cat#sc-58774) were from Santa Cruz. PDIA6 antibody (cat#18233-1-AP) was from ProteinTech. UGGT1 antibody (cat#ADI-VAP-PT068) was from Enzo Life Sciences. Human Cpep-specific (Mouse monoclonal, 20G11) antibody was kindly provided by Dr. Bill Balch, Scripps Res. Inst. BiP rabbit polyclonal antibody was a kind gift from Dr. Linda Hendershot (St. Jude Children's Research Hospital, Memphis, TN). Flag antibody (cat#F1804-200UG) was purchased from Sigma Aldrich. SEL1L antibody (cat#ab78298) was obtained from Abcam. FVIII antibody (cat#GMA012) was obtained from Green Mountain Antibodies (Burlington, VT). FVIII heavy chain monoclonal antibody F8 coupled to Sepharose CL-4B was a kind gift from Baxter/Baxalta Corp. CNX rabbit polyclonal antibody was kindly provided by Dr. Maurizio Molinari. Gold-labelled secondary murine antibodies were kindly provided by the UCSD Microscopy Core. Prolong Antifade Gold murine FAB fragments, Dylight 549 conjugated anti-mouse FAB fragments and Texas-Red conjugated anti-mouse secondary were obtained from Jackson Immunoresearch.

##### Cell lines and transfection.

Parental Chinese Hamster Ovary cells (CHO-K1), CHO cells that constitutively express FVIII, clones 10A1[1] and H9[2], were previously described. CHO cells were grown in α-MEM (Corning®, Life Sciences) (containing 5mM glucose, 2.5mM L-glutamine) supplemented with 10% FBS and 1% Penicillin/Streptomycin. H9 and 10A1 cells were propagated in 0.1μM methotrexate (mtx). COS-1 and 293T cells were grown in DMEM (Corning®, Life Sciences) (containing 4.5g/L D-glucose, 2.5mM L-glutamine) supplemented with 10% FBS, 110mg/L sodium pyruvate and 1% Penicillin/Streptomycin. Transient transfection was performed using Lipofectamine 3000 (Invitrogen) following manufacturer's instructions.

##### Plasmid construction.

To construct wtFVIII-CpepMyc (wtFVIII-CMy), 79aa (residues 253-331) from the FVIII A1 domain were amplified from pSFFV-wtFVIII[3] by Phusion DNA polymerase (ThermoFisher Scientific) using primers: FVIIIApaI forward: 5'- TAGGGCCCAGTCTATTGGCATGTGATTGG-3' and FVIIIApaI reverse: 5'- ATGGGCCCGAGATATGACAAAACAGTAGAAAC-3'. The 79aa PCR fragment was digested with *Apa*I and ligated into the proinsulin expression vector (pTarget-hProCpepMyc)[4] at the *Apa*I cloning site to yield pTarget-wtFVIII-hProCpepMyc. Then the 366bp of wtFVIII-hCpepMyc coding sequence was amplified from pTarget-wtFVIII-hProCpepMyc using FVIII-CpepPmeI forward: 5'-TCTGTTTAAACGGGAGGCAGAGGACCTGCAGG-3' and FVIII-CpepXbaI reverse: 5'-GATTCTAGACTACTGCAGGGACCCCTCCAGGGC-3' primers and subsequently ligated into the *Pme*I and *Xba*I cloning sites in pTarget vector to generate pTarget-wtFVIII-CMy. The F309SFVIII-CMy, FV-CMy and eGFP-CMy were generated similarly as wtFVIII-CMy. wtFVIII- and mutants-eGFP-CMy were constructed similarly as FVIII-CMy by using pTarget-eGFP-CMy as template. The cDNA sequences were all verified by restriction enzyme digestion and DNA sequencing. All FVIII expression plasmids[5], and pMT2-B-domain-deleted FVIII (BDD-FVIII)[6] were previously described. BiP expression plasmids (BiP-Flag, V461FBiP-Flag and BiP-M)[7] were a gift from Dr. David Ron (Cambridge Institute for Medical Research, UK).

**Metabolic inhibition.**

ATP depletion/repletion experiments were performed in induced H9-CHO cells and transfected 293T cells as previous described[8].

**Radiolabeling and IP.**

10A1 CHO cells were pulse-labeled in 0.5ml Met/Cys-free DMEM containing [$^{35}$S]-Met/Cys (300µCi/mL) for 20min. Then, chase was performed for the indicated times in medium with or without 20mM 2DG and 10mM $NaN_3$ containing excess unlabeled (5mM) (Met/Cys) and 10mg/ml aprotinin. For repletion, metabolic inhibitors were removed and replaced with regular medium containing Glc, excess unlabeled 5mM Cys/Met, and aprotinin as above. At the end of the chase period, conditioned media were collected and cell lysates prepared in lysis buffer [50mM Tris-HCl, pH 7.4, 150mM NaCl, 0.1%(v/v) Triton X-100, and 1%(v/v) IGEPAL] containing protease inhibitor cocktail and 1mM phenylmethyl sulfonyl fluoride (PMSF). Equal amounts of cell lysates and corresponding amounts of media were subjected to FVIII immunoprecipitation (IP) using FVIII antibody coupled to Sepharose CL-4B beads (Baxter Corp, Deerfield, IL). IP'ed FVIII was separated by SDS-PAGE on a 6% polyacrylamide gel and visualized by autoradiography.

**FVIII activity measurement.**

FVIII activity was measured by the 1-stage activated partial thromboplastin time (aPTT) clotting assay on an MLA Electra 750 fibrinometer (Medical Laboratory Automation, Pleasantville, NY) by reconstitution of human FVIII-deficient plasma using FACT plasma (normal pooled plasma from George King Biomedical) as a standard. FVIII antigen was quantified by a FVIII sandwich ELISA using the Affinity Biologicals FVIII:C-EIA kit following manufacturers' instructions.

**Fluorescence microscopy.**

H9 cells were treated as in Fig. 1B legend and then fixed in 4% paraformaldehyde (PFA) for 10min and permeabilized with PBS containing 0.5% Triton X-100 and 1% BSA for 5min. The permeabilized cells were incubated with thioflavin-S (0.05% w/v) in 70% ethanol for 8min. Cells were sequentially washed with 70% ethanol for 10min and washed with water for another 10min before blocking and probed with indicated antibodies (PDIA6, KDEL, and FVIII). Finally, cells were incubated with Texas-Red conjugated mouse secondary for 1h at RT, followed by 3 washes with blocking buffer and 3 washes with PBS. Coverslips were applied with Prolong Antifade Gold.

**Sucrose gradient velocity sedimentation**.

Cell lysates from NaB-treated H9 CHO cells or transfected 293T cells were prepared as previously described[8]. After centrifugation, protein from each fraction (1-10) was collected and analyzed by IP, Western blotting, TEM, cryo-EM or subjected to filtration assay.

**Filtration assay.**

Cell lysates were applied to either 0.22µm cellulose acetate (CA) or nitrocellulose (NC) (Pall Corp, FL) membranes pre-equilibrated with PBS on a Bio-Dot microfiltration apparatus (Bio-Rad) and subjected to vacuum filtration as described[9]. Membranes were subjected to Western blotting using specified antibodies. Densities were quantified by ECL or Li-Cor imaging systems.

**Immunogold labeling TEM.**

CHO-K1 or FVIII treated with or without SAHA (5µM) for 24h and then fixed in 4% paraformaldehyde in 0.1M PBS for 5min without washing and left at 4$^{o}$C overnight. Further processing was performed by the Microscopy Core at the University of California, San Diego (UCSD). Images were obtained using a Tecnai G2 Spirit BioTWINTEM equipped with an Eagle 4k HS digital camera (FEI, Hilsboro, OR) with indicated magnifications.

**Negative stain TEM.**

Lysates from CHO-K1 or H9 cells treated with NaB for 24h were applied to 5-20% sucrose gradients for velocity sedimentation. FVIII in each fraction was IP'ed with monoclonal antibody conjugated Sepharose beads and eluted by incubation with 50% ethylene glycol at 4$^{o}$C for 1h. Aliquots of IP'ed FVIII were added to carbon grids for 5min, prior to staining for 5min with 1%

phosphotungstic acid. After drying, the screens were visualized at a magnification from 23,000-92,000X using a JEOL JSM 1400 transmission electron microscope in the University of Michigan Microscopy Lab.

**Transmission electron cryo-microscopy (cryo-EM) and analysis.**

H9 cells were treated with SAHA (5μM) for 24h and lysates prepared for sucrose gradient sedimentation. FVIII was IP'ed from heavy and light sucrose fractions in glycine buffers pH 4.5/4/3 for the cryo-EM studies. Sepharose beads stored in PBS were transferred to glycine buffer (100mM, pH 4.5), resuspended, and incubated at RT followed by a short 37$^{o}$ C incubation. A drop of 5μl sample was applied to plasma-cleaned (Solaris, Gatan Inc.) Quantifoil 1.2/1.3 holey carbon-coated 200 mesh electron microscopy grids (Quantifoil Micro Tools GmbH). Excess liquid was blotted, and the grids were then manually frozen in liquid N2-cooled ethane. Images of all samples suspended over holes were acquired under low-dose conditions using TecnaiT12 G2 Twin TEM microscopes (Thermo Fisher, FEI company) operated at 120 kV (Lab6) at nominal magnification of 52,000X. Immuno-Gold labeling of purified FVIII fractions from NaB treated CHO and H9 cells was performed with human FVIII (heavy chain) monoclonal antibody (Baxter Corp, Deerfield, IL) as primary and a 10-nm gold-conjugated anti-mouse IgG (goat, affinity isolated; Sigma-Aldrich G7652) as secondary antibodies. Controls included (i) exclusion of primary antibody and (ii) uninduced H9 cells. Processing was done in pyCoAn, an extended python-version of CoAn[10] and in EMAN2[11]. Micrographs were corrected for background variations. Overlapping boxes were selected along the path of straight, single fibril segments and aligned with the fibril axis. Boxes were subjected to k-means clustering with approximately 50 boxes per class. Class averages were calculated and projected down the fibril axis to form a 1-dimensional profile. Diameters for each class were determined using the inflection points of the 1-dimensional profiles.

**Affinity purification mass spectrometry (AP-MS).**

CHO-K1 and H9 cells were plated onto 4x15 cm dishes and treated with 5mM NaB for 21h. The cells were then lysed in 50mM Tris-HCl, 150mM NaCl, 2% (w/v) CHAPS containing buffer with protease inhibitors on ice for 40min. Lysates were centrifuged at 12,000xg for 15min. One mg of protein was precleared with Protein A beads for 1h at 4$^{o}$C. The supernatants were incubated with 400µl of Baxter FVIII-CL-4B beads (50% slurry) overnight at 4$^{o}$C. Beads were washed twice with detergent containing buffer and eluted with 50mM Tris-HCl, 1%SDS at 95$^{o}$C for 10min. Samples were analyzed by 2D liquid chromatography coupled to tandem mass spectrometry (2DLC-MS/MS). Briefly, proteins were reduced with tris(2-carboxyethyl)phosphine (TCEP), alkylated with iodoacetamide (IAA), and subjected to overnight digestion with mass spec grade Trypsin/Lys-C mix (Promega, Madison, WI). Peptides were then analyzed by 2DLC-MS/MS using a 2D nanoACQUITY Ultra Performance Liquid Chromatography (UPLC) system (Waters corp., Milford, MA) coupled to a Q-Exactive Plus mass spectrometer (Thermo Fisher Scientific). Peptides were loaded onto the first-dimension column, XBridge BEH130 $C_{18}$NanoEase (300µm x 50mm, 5µm) equilibrated with solvent A (20mM ammonium formate pH 10, first dimension pump) at 2µL/min. The first fraction was eluted from the first dimension column at 17% of solvent B (100% acetonitrile) for 4min and transferred to the second dimension Symmetry C18 trap column 0.180 x 20 mm (Waters corp., Milford, MA) using a 1:10 dilution with 99.9% second dimensional pump solvent A (0.1% formic acid in water) at 20µL/min. Peptides were then eluted from the trap column and resolved on the analytical $C_{18}$ BEH130 PicoChip column 0.075 x 100 mm, 1.7µm particles (New Objective, MA) at low pH by increasing the composition of solvent B (100% acetonitrile) from 2 to 26% over 94min at 400 nL/min. Subsequent fractions were treated with increasing concentrations of solvent B. The following 4 first dimension fractions were eluted at 19.5, 22, 26, and 65% solvent B. The mass spectrometer was operated in positive data-dependent acquisition mode. MS1 spectra were measured with a resolution of 70,000, an AGC target of 1e6 and a mass range from 350 to 1700 m/z. Up to 12 MS2 spectra per duty cycle were triggered, fragmented by HCD, and acquired with a resolution of 17,500 and an AGC target of 5e4, an isolation window of 2.0 m/z and a normalized collision energy of 25. Dynamic exclusion was enabled with duration of 20 sec. All mass spectra were analyzed with MaxQuant software version 1.5.2.8. MS/MS spectra were searched against the *Homo sapiens* Uniprot protein sequence database (version January 2016) and GPM cRAP sequences (commonly known protein contaminants). Precursor mass tolerance was set to 20ppm and 4.5ppm for the first search where initial mass recalibration was completed and for the main search, respectively. Product ions were searched with a mass tolerance 0.5 Da. The maximum precursor ion charge state used for searching was 7. Carbamidomethylation of cysteines was searched as a fixed modification, while methionine oxidation and acetylation of protein N-terminal were searched as variable modifications. Enzyme was set to trypsin in a specific mode and a maximum of two missed cleavages was allowed for searching. The target-decoy-based false discovery rate (FDR) filter for spectrum and protein identification was set to 1%.

**Co-Immunoprecipitations (Co-IPs).**

Co-IPs were performed using the cell lysates from either H9 cells or transfected 293T cells. Protein extracts ~200µg were incubated with indicated antibodies (Cpep or Flag) in lysis buffer (150mM NaCl, 50mM Tris [pH 7.4]) supplemented with protease inhibitors (Roche) for 16h at

4ºC. Antigen-antibody complexes were then incubated with protein A/G sepharose beads (BioVision) for 4h at 4ºC. The beads were washed 3x with 500µl lysis buffer and complexes were eluted by boiling at 95ºC for 10min in 2x sample buffer. Eluted proteins were analyzed by Western blotting with indicated antibodies. FVIII IP was performed as described above by using FVIII antibody conjugated sepharose beads.

**References**


1. Kaufman RJ, Wasley LC, Davies MV, Wise RJ, Israel DI, Dorner AJ. Effect of von willebrand factor coexpression on the synthesis and secretion of factor viii in chinese hamster ovary cells. *Mol Cell Biol*. 1989;9:1233-1242.
2. Dorner AJ, Wasley LC, Kaufman RJ. Increased synthesis of secreted proteins induces expression of glucose-regulated proteins in butyrate-treated chinese hamster ovary cells. *J Biol Chem*. 1989;264:20602-20607.
3. Ward NJ, Buckley SMK, Waddington SN, VandenDriessche T, Chuah MKL, Nathwani AC, McIntosh J, Tuddenham EGD, Kinnon C, Thrasher AJ, McVey JH. Codon optimization of human factor viii cdnas leads to high-level expression. *Blood*. 2011;117:798-807.
4. Liu M, Haataja L, Wright J, Wickramasinghe NP, Hua Q-X, Phillips NF, Barbetti F, Weiss MA, Arvan P. Mutant ins-gene induced diabetes of youth: Proinsulin cysteine residues impose dominant-negative inhibition on wild-type proinsulin transport. *PLoS One*. 2010;5:e13333-e13333.
5. Swaroop M, Moussalli M, Pipe SW, Kaufman RJ. Mutagenesis of a potential immunoglobulin-binding protein-binding site enhances secretion of coagulation factor viii. *J Biol Chem*. 1997;272:24121-24124.
6. Gilbert GE, Kaufman RJ, Arena AA, Miao H, Pipe SW. Four hydrophobic amino acids of the factor viii c2 domain are constituents of both the membrane-binding and von willebrand factor-binding motifs. *J Biol Chem*. 2002;277:6374-6381.
7. Preissler S, Chambers JE, Crespillo-Casado A, Avezov E, Miranda E, Perez J, Hendershot LM, Harding HP, Ron D. Physiological modulation of bip activity by trans-protomer engagement of the interdomain linker. *eLife*. 2015;4:e08961.
8. Tagliavacca L, Wang Q, Kaufman RJ. Atp-dependent dissociation of non-disulfide-linked aggregates of coagulation factor viii is a rate-limiting step for secretion. *Biochemistry*. 2000;39:1973-1981.

9. Wanker EE, Scherzinger E, Heiser V, Sittler A, Eickhoff H, Lehrach H. Membrane filter assay for detection of amyloid-like polyglutamine-containing protein aggregates. *Methods enzymol*. Academic Press; 1999:375-386.
10. Volkmann N, Hanein D. Quantitative fitting of atomic models into observed densities derived by electron microscopy. *J Struct Biol*. 1999;125:176-184.
11. Tang G, Peng L, Baldwin PR, Mann DS, Jiang W, Rees I, Ludtke SJ. Eman2: An extensible image processing suite for electron microscopy. *J Struct Biol*. 2007;157:38-46.

**Supplemental figure legends**

**Figure S1. Metabolic collapse causes FVIII amyloid-like aggregation.** 10A1 cells treated without (i) or with 2DG+$NaN_3$ for 2h (ii) and subsequently recovered in Glc-containing media for 4h (iii) were stained the β-sheet binding dye Thio-S (green) and with antibodies for FVIII (red) and KDEL (blue). The merged images demonstrate co-localization of FVIII, KDEL, and Thio-S.

**Figure S2. Glc metabolism, not oxidative phosphorylation, is required for FVIII solubility. A.** FVIII binds CA membranes. FVIII aggregation was monitored by CA and NC filtration of lysates from H9 cells treated with or without NaB for 24h or from 10A1 cells. Indicated cells were untreated or treated with Glc-free media containing 20mM 2DG and 10mM $NaN_3$ for 2h. Recombinant FVIII (rFVIII) was filtered as control for well-folded soluble FVIII. Membranes were probed with FVIII antibody. **B.** ATP is depleted upon 2DG + $NaN_3$ treatment**.** H9 cells were treated with NaB for 17h or with 20mM 2DG+10mM $NaN_3$ for 2h. Uninduced, induced and metabolically inhibited cells were trypsinized for quantification of ATP by the ATPlite kit (PerkinElmer) according to the manufacturer's protocol. The luciferase signal was recorded on a Veritas Microplate Luminometer (Turner BioSystem). Triplicate ATP measurements were analyzed with one-way ANOVA provided by GraphPad Prism software.****: $p<0.0001$ compared to untreated cells, ns = non-statistically significant. Results are presented as ATP/cell. **C**. FVIII aggregates are not degraded via proteasomal or autophagy pathways**.** H9 cells treated with NaB for 24h and 10A1 cells were both treated with 2DG+$NaN_3$ followed by recovery in Glc-containing medium in the absence or presence of proteasome inhibitor (Velcade, Vel) or autophagy inhibitor (3-methyladenine, 3MA) for 4h. Cell lysates were analyzed for FVIII aggregation by CA membrane filtration. **D**. Oxidative phosphorylation is not required to prevent FVIII aggregation**.** H9 cells were treated with 5mM NaB for 18h and then treated with 1μM oligomycin A (Oligo), a complex V inhibitor, in complete α-MEM or with 20mM 2DG+10mM $NaN_3$ in DMEM medium without Glc for 2h. For repletion, 2DG+$NaN_3$ containing medium was replaced with complete α-MEM and cells were cultured for an additional 4h. FVIII aggregation from metabolically inhibited cells was compared to untreated (UT) cells. CA membranes were probed with β-actin antibody as a loading control.

**Figure S3. A.** Schematic represents the wtFVIII-CMy and wtFVIII-eGFP-CMy chimeras. 79aa from the FVIII A1 domain (residues 253- 331) were inserted in frame upstream of the human proinsulin C peptide-Myc (CMy). cMyc-epitope tag is highlighted in box, human C-peptide sequences are underlined. SP is the proinsulin signal peptide. The amino acids corresponding to copper ion ligands (aa His267/Cys310/His315) are depicted in red bold. Phe309 relative to the first aa of mature FVIII is indicated in red. Cys329 was mutated to Gly to prevent aberrant disulfide bond formation. N-liked glycosylation sites (N-Gly) and disulfide bonds (S-S) in full-length FVIII are indicated**. B.** wtFVIII-CMy is expressed in 293T cells. Cell lysates from wtFVIII-CMy transfected 293T cells were analyzed by Western blotting using Cpep and vinculin antibodies. Cell lysates and culture media from CMy, F309SFVIII-CMy, FV-CMy and eGFP-CMy transfected cells were included as controls.

**Figure S4. Prediction of β-aggregate propensity in the 79aa of the FVIII A1 domain.** The β-aggregate propensity scores were calculated using the TANGO algorithm[1], for the 79aa of the A1 domain: (**A**) wtFVIII, (**B**) F306W, (**C**) F309S, (**D**) C310S and (**E**) C310E. In addition, the aggregation propensity for the homologous region of FV was calculated (**F**). Histograms display aggregation propensity vs the linear aa sequence.

**Figure S5. A 79aa motif independently forms high molecular weight (HMW) aggregates.** wtFVIII-Cpep aggregates into HMW complexes. Lysates from 293T cells expressing wtFVIII-CMy, F309SFVIII-CMy or eGFP-CMy were subjected to sucrose gradient sedimentation as in Fig. 3G. Protein samples from each fraction (1-10) were analyzed by Western blotting using Cpep antibody. The membrane was probed for β-actin for protein loading control. Sucrose gradient of cell lysate from 293T cells transfected with a full-length wtFVIII expression vector was included as a control. wtFVIII exhibits very HMW aggregates. The percentage of protein in each fraction is indicated. Red arrows indicate migration of soluble forms of F309SFVIII-CMy.

**Figure S6. Affinity Purification-Mass Spectrometry (AP-MS) identifies proteins that interact with FVIII.** H9 cells treated with NaB for 21h were IP'ed with FVIII antibody conjugated to Sepharose beads. The parental CHO cells were included as a negative control. **A.** Silver staining of IP'ed FVIII from biological triplicates. Migration of FVIII and BiP are indicated. LC and HC indicate IgG light and heavy chains. **B**. ER chaperones are enriched in IP'ed FVIII. Heat map of ER resident proteins detected in the IP'ed FVIII from NaB-treated H9 cells showing spectral counts in each sample replicate. ER resident proteins are clustered according to their protein family. **C**. For validation, equal percentages of samples from total cell lysates, IP'ed FVIII and the post-IP supernatants were analyzed by reducing SDS-PAGE for blotting with the indicated antibodies. ERP44 was included as a negative control. ERGIC-53 was included as another negative control for a weak interactor with FVIII[2] but not detected in the AP-MS analysis.

**Figure S7. Calnexin/Calreticulin (CANX/CRT) prevent FVIII aggregation. A**. Figure depicts the action of the α-glucosidase inhibitor castanospermine (CST) on GS1 and GS2 in preventing entry into the CANX/CRT cycle. The red circle indicates the glucose residues that are trimmed by GS1 and GS2. **B**. H9 cells were treated with SAHA for 18h and then treated with increasing concentrations of CST from 5 to 500 µM for 6h. Cell lysates were prepared for analysis by filtration (B) and Western blotting (C) using indicated antibodies. Cells treated with 10µg/ml Tm were included for a positive control for FVIII aggregation. Levels of FVIII aggregation were quantified from FVIII signals on CA membranes relative to the Western blot. Cells treated with vehicle DMSO (V) were used as a negative control. CST increases FVIII aggregation at ~100µM, which is the concentration required to inhibit GS1 and GS2 in cells[3,4]. **D-E**. wtFVIII-CMy aggregation is glycosylation independent. 293T cells expressing wtFVIII-CMy were treated with 200µM CST for 6h and cell lysates were prepared for filtration on CA (**D**) and Western blotting with indicated antibodies (**E**). Cells transfected with eGFP-CMy and FV-CMy were included as controls. Treating the cells with 10µg/ml Tm was included as positive control for protein aggregation. Cpep aggregation under CST and Tm treatment was quantified as percentage relative to the untreated condition. The FV sequence has one N-linked glycan as exhibits increased aggregation upon CST treatment. *, ** represent non-glucose trimmed and non-glycosylated FV-CMy, respectively.

**Figure S8. Forced BiP expression prevents FVIII aggregation.** 293T cells were transfected with expression vectors for BDD-FVIII and BiP-Flag or V461FBiP-Flag (peptide-binding defective BiP). For the last 2.5h, cells were treated with or without 2DG+$NaN_3$. FVIII aggregation in cell lysates was measured by Western blotting (**A**) and CA filtration (**B**). BiP-Myc (BiP-M) was included as a biological replicate for the impact of BiP on FVIII aggregation.

**Figure S9. BiP depletion by SubAB protease does not affect eGFP-CMy secretion.** 293T cells expressing eGFP-CMy were treated with 2µg/ml SubAB protease for 2.5h. Then cell lysates and cultured media were analyzed by Western blotting for BiP cleavage (**A**) and eGFP-CMy secretion (**B**). Data show biological replicates.

#### References


1. Fernandez-Escamilla AM, Rousseau F, Schymkowitz J, Serrano L. Prediction of sequence-dependent and mutational effects on the aggregation of peptides and proteins. *Nat Biotechnol*. 2004;22:1302-1306.
2. Zhang B, Kaufman RJ, Ginsburg D. Lman1 and mcfd2 form a cargo receptor complex and interact with coagulation factor viii in the early secretory pathway. *J Biol Chem*. 2005;280:25881-25886.
3. Kaushal GP, Pan YT, Tropea JE, Mitchell M, Liu P, Elbein AD. Selective inhibition of glycoprotein-processing enzymes. Differential inhibition of glucosidases i and ii in cell culture. *J Biol Chem*. 1988;263:17278-17283.

4. Taylor DL, Kang MS, Brennan TM, Bridges CG, Sunkara PS, Tyms AS. Inhibition of alpha-glucosidase i of the glycoprotein-processing enzymes by 6-o-butanoyl castanospermine (mdl 28,574) and its consequences in human immunodeficiency virus-infected t cells. *Antimicrob Agents Chemother*. 1994;38:1780-1787.

### Figure S1

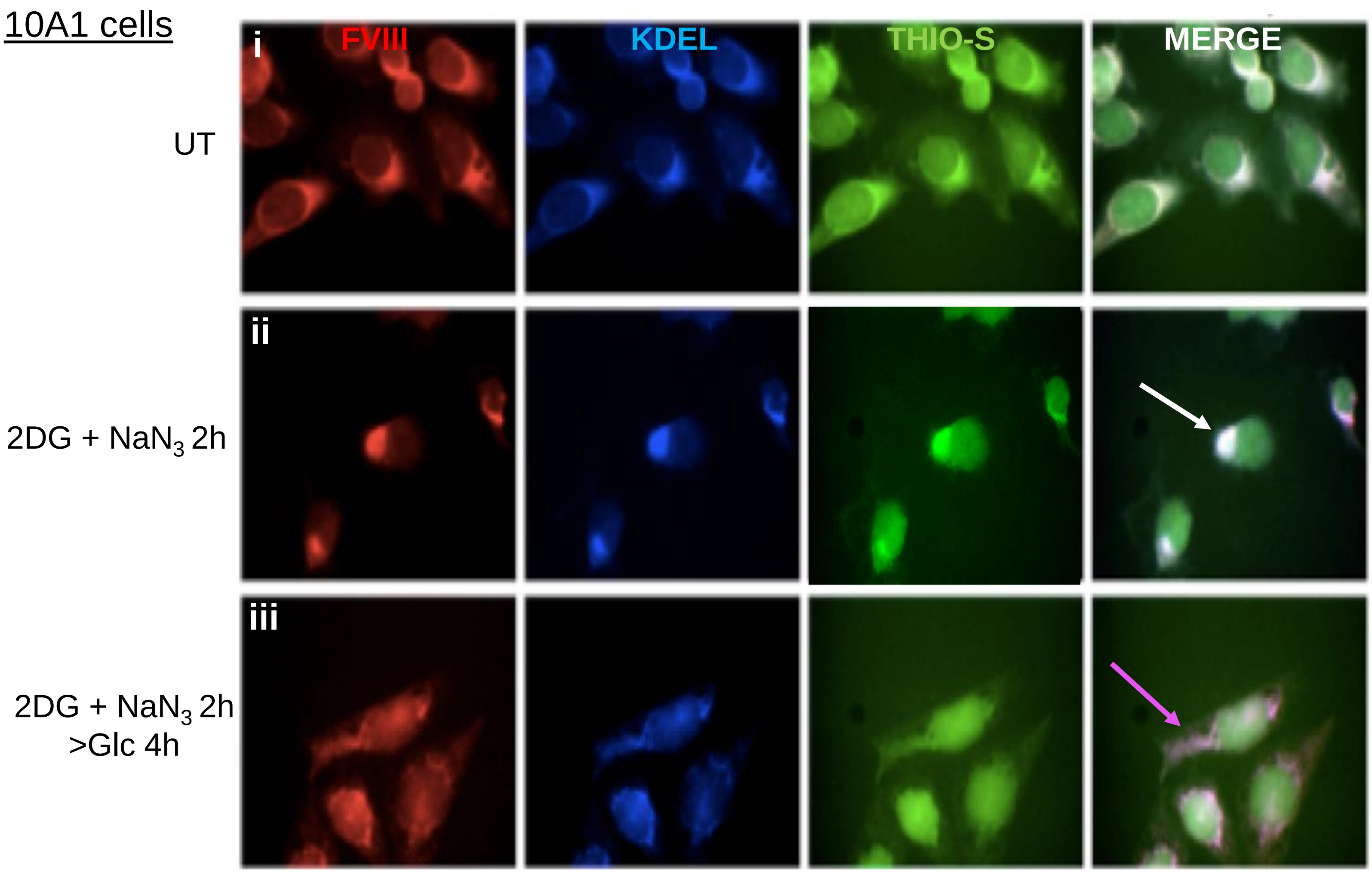
10A1 cells
i
FVIII
KDEL
THIO-S
MERGE
UT
ii
2DG + NaN3 2h
iii
2DG + NaN3 2h
>Glc 4h

Figure S2

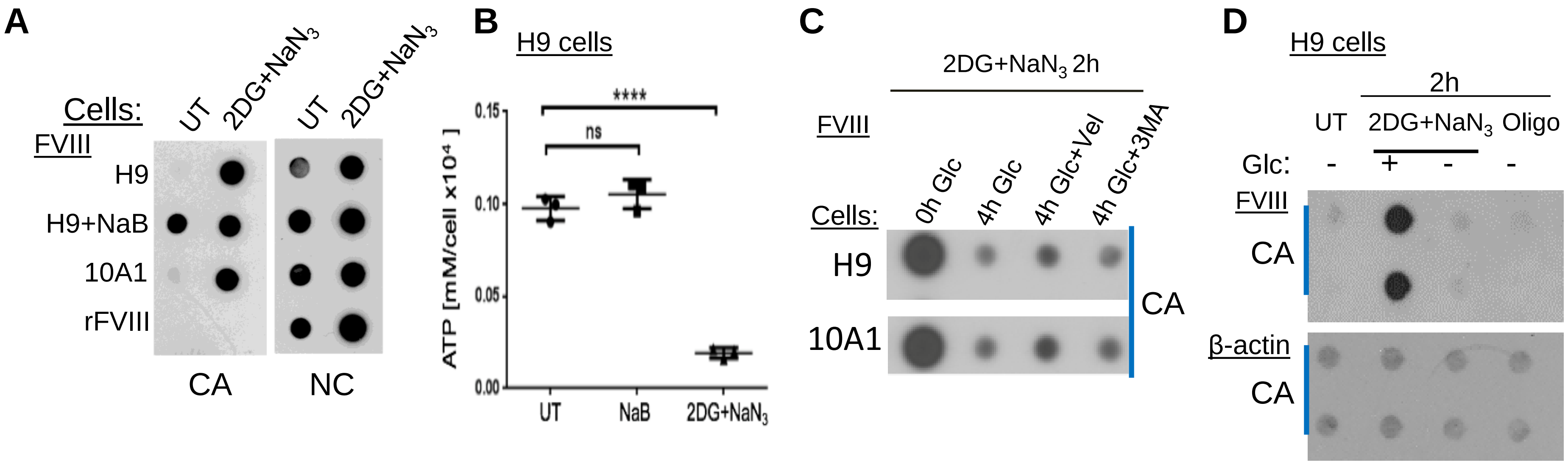

A
Cells:
UT
2DG+NaN3
UT
2DG+NaN3
FVIII
H9
H9+NaB
10A1
rFVIII
CA
NC
B
H9 cells
ATP [mM/cell x10⁴ ]
0.15
0.10
0.05
0.00
****
ns
UT
NaB
2DG+NaN3
C
2DG+NaN3 2h
FVIII
0h Glc
4h Glc
4h Glc+Vel
4h Glc+3MA
Cells:
H9
10A1
CA
D
H9 cells
2h
UT
2DG+NaN3
Oligo
Glc:
-
+
-
-
FVIII
CA
β-actin
CA

Figure S3

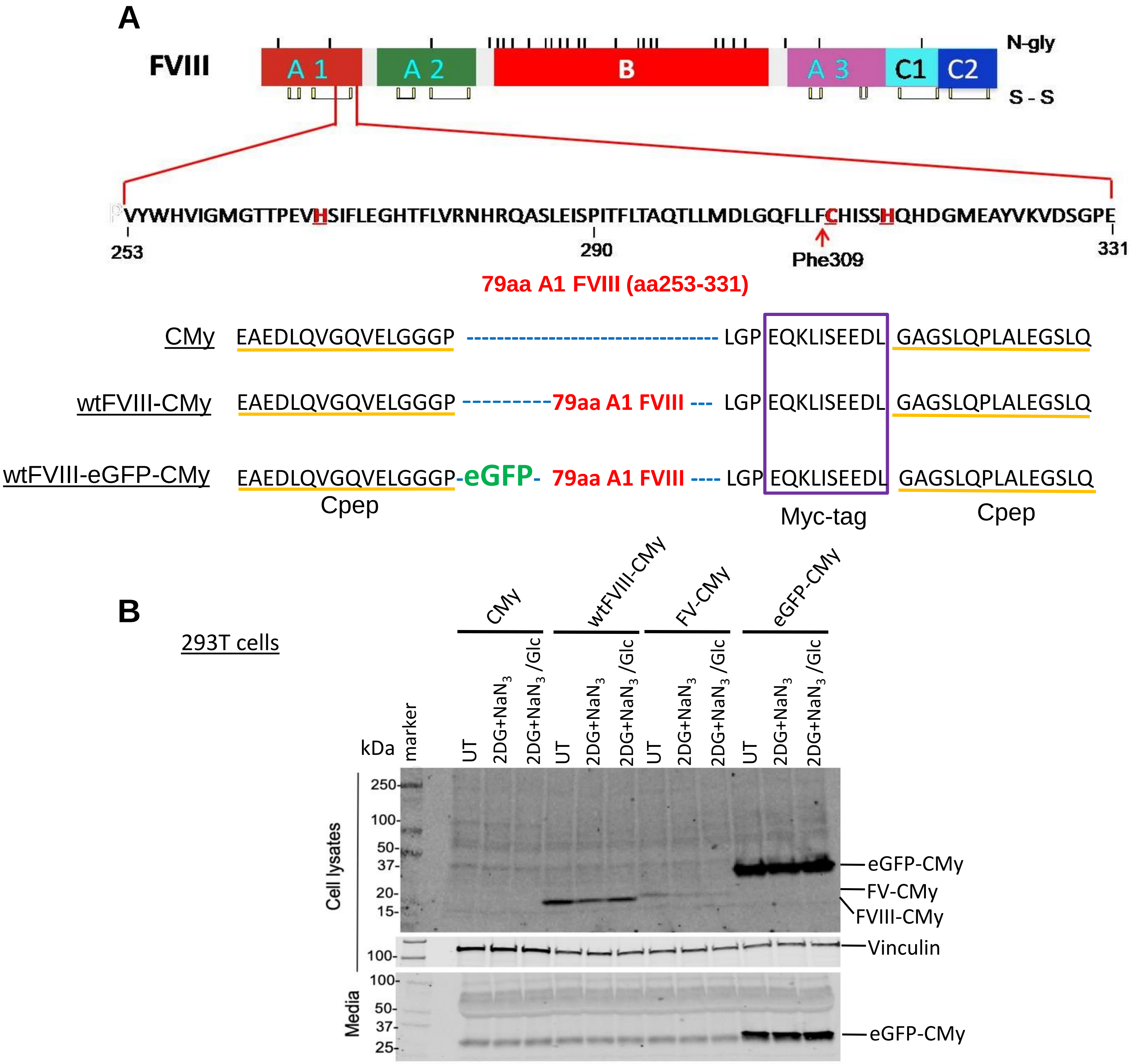
A
FVIII
N-gly
A 1
A 2
B
A 3
C1
C2
S - S
VYWHVIGMGTTPEVHSIFLEGHTFLVRNHRQASLEISPITFLTAQTLLMDLGQFLLFCHISSHQHDGMEAYVKVDSGPE
253
290
Phe309
331
79aa A1 FVIII (aa253-331)
CMy EAEDLQVGQVELGGGP LGP EQKLISEEDL GAGSLQPLALEGSLQ
wtFVIII-CMy EAEDLQVGQVELGGGP 79aa A1 FVIII LGP EQKLISEEDL GAGSLQPLALEGSLQ
wtFVIII-eGFP-CMy EAEDLQVGQVELGGGP-eGFP- 79aa A1 FVIII LGP EQKLISEEDL GAGSLQPLALEGSLQ
Cpep
Myc-tag
Cpep
B
293T cells
CMy
wtFVIII-CMy
FV-CMy
eGFP-CMy
kDa
marker
UT
2DG+NaN3
2DG+NaN3 /Glc
Cell lysates
Media
250-
100-
50-
37-
20-
15-
100-
100-
50-
37-
25-
eGFP-CMy
FV-CMy
FVIII-CMy
Vinculin
eGFP-CMy

Figure S4

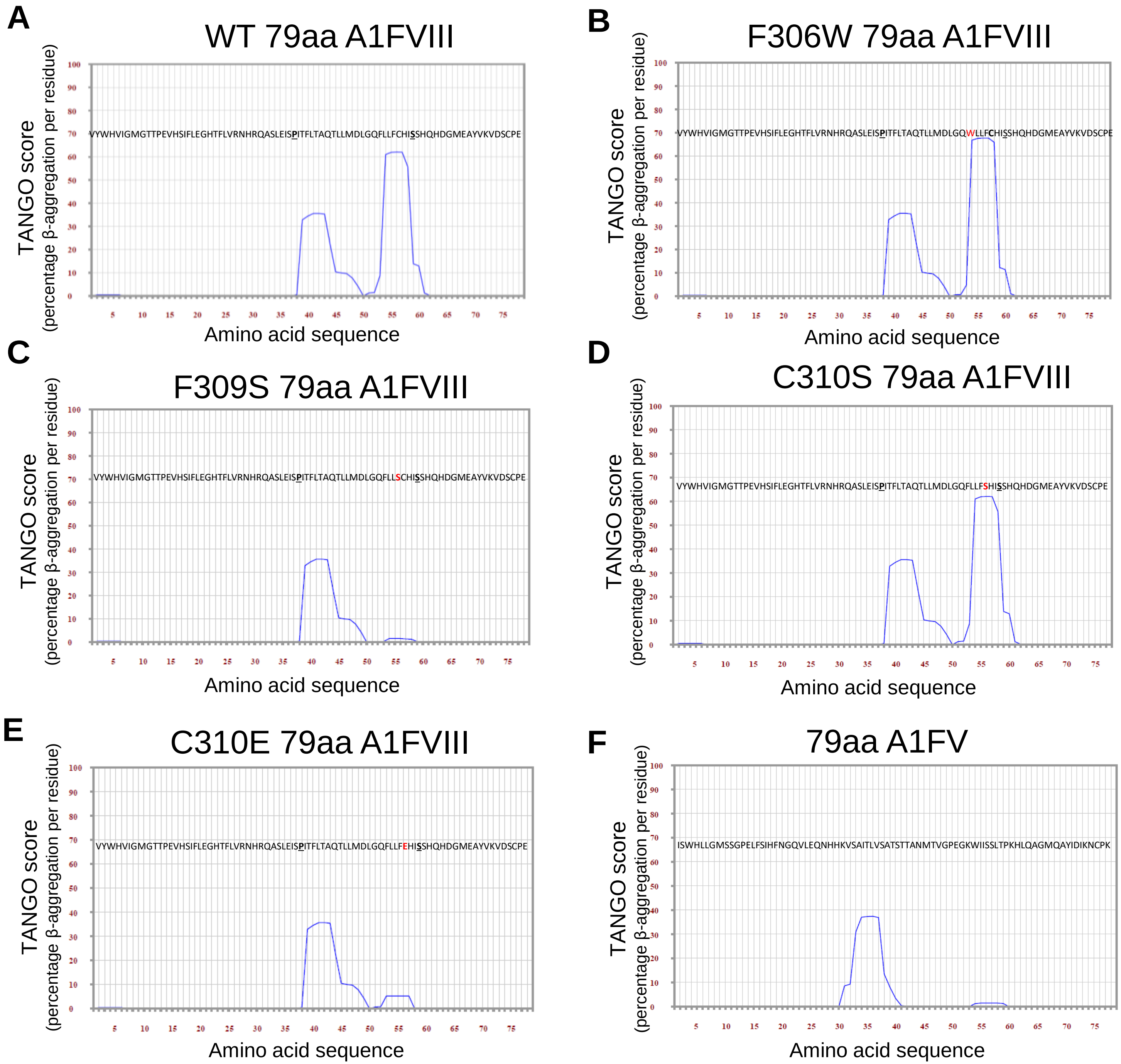
A
WT 79aa A1FVIII
VYWHVIGMGTTPEVHSIFLEGHTFLVRNHRQASLEISPITFLTAQTLLMDLGQFLLFCHISSHQHDGMEAYVKVDSCPE
B
F306W 79aa A1FVIII
VYWHVIGMGTTPEVHSIFLEGHTFLVRNHRQASLEISPITFLTAQTLLMDLGQWLLFCHISSHQHDGMEAYVKVDSCPE
C
F309S 79aa A1FVIII
VYWHVIGMGTTPEVHSIFLEGHTFLVRNHRQASLEISPITFLTAQTLLMDLGQFLLSCHISSHQHDGMEAYVKVDSCPE
D
C310S 79aa A1FVIII
VYWHVIGMGTTPEVHSIFLEGHTFLVRNHRQASLEISPITFLTAQTLLMDLGQFLLFSHISSHQHDGMEAYVKVDSCPE
E
C310E 79aa A1FVIII
VYWHVIGMGTTPEVHSIFLEGHTFLVRNHRQASLEISPITFLTAQTLLMDLGQFLLFEHISSHQHDGMEAYVKVDSCPE
F
79aa A1FV
ISWHLLGMSSGPELFSIHFNGQVLEQNHHKVSAITLVSATSTTANMTVGPEGKWIISSLTPKHLQAGMQAYIDIKNCPK
TANGO score
(percentage β-aggregation per residue)
Amino acid sequence

### Figure S5

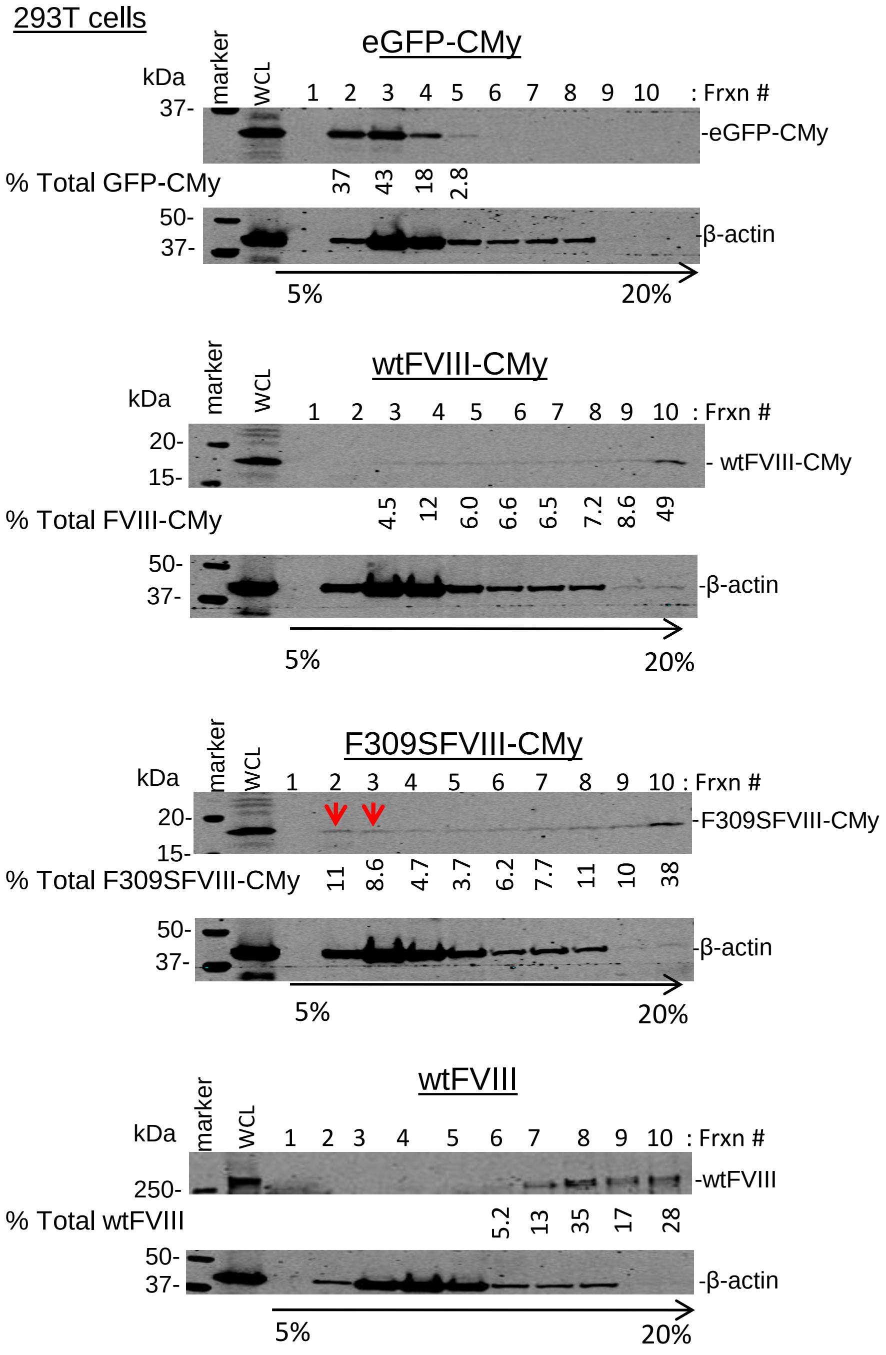
293T cells
eGFP-CMy
kDa
marker
WCL
1 2 3 4 5 6 7 8 9 10 : Frxn #
37-
-eGFP-CMy
% Total GFP-CMy
37 43 18 2.8
50-
37-
-β-actin
5%
20%
wtFVIII-CMy
kDa
marker
WCL
1 2 3 4 5 6 7 8 9 10 : Frxn #
20-
15-
- wtFVIII-CMy
% Total FVIII-CMy
4.5 12 6.0 6.6 6.5 7.2 8.6 49
50-
37-
-β-actin
5%
20%
F309SFVIII-CMy
kDa
marker
WCL
1 2 3 4 5 6 7 8 9 10 : Frxn #
20-
15-
-F309SFVIII-CMy
% Total F309SFVIII-CMy
11 8.6 4.7 3.7 6.2 7.7 11 10 38
50-
37-
-β-actin
5%
20%
wtFVIII
kDa
marker
WCL
1 2 3 4 5 6 7 8 9 10 : Frxn #
250-
-wtFVIII
% Total wtFVIII
5.2 13 35 17 28
50-
37-
-β-actin
5%
20%

Figure S6

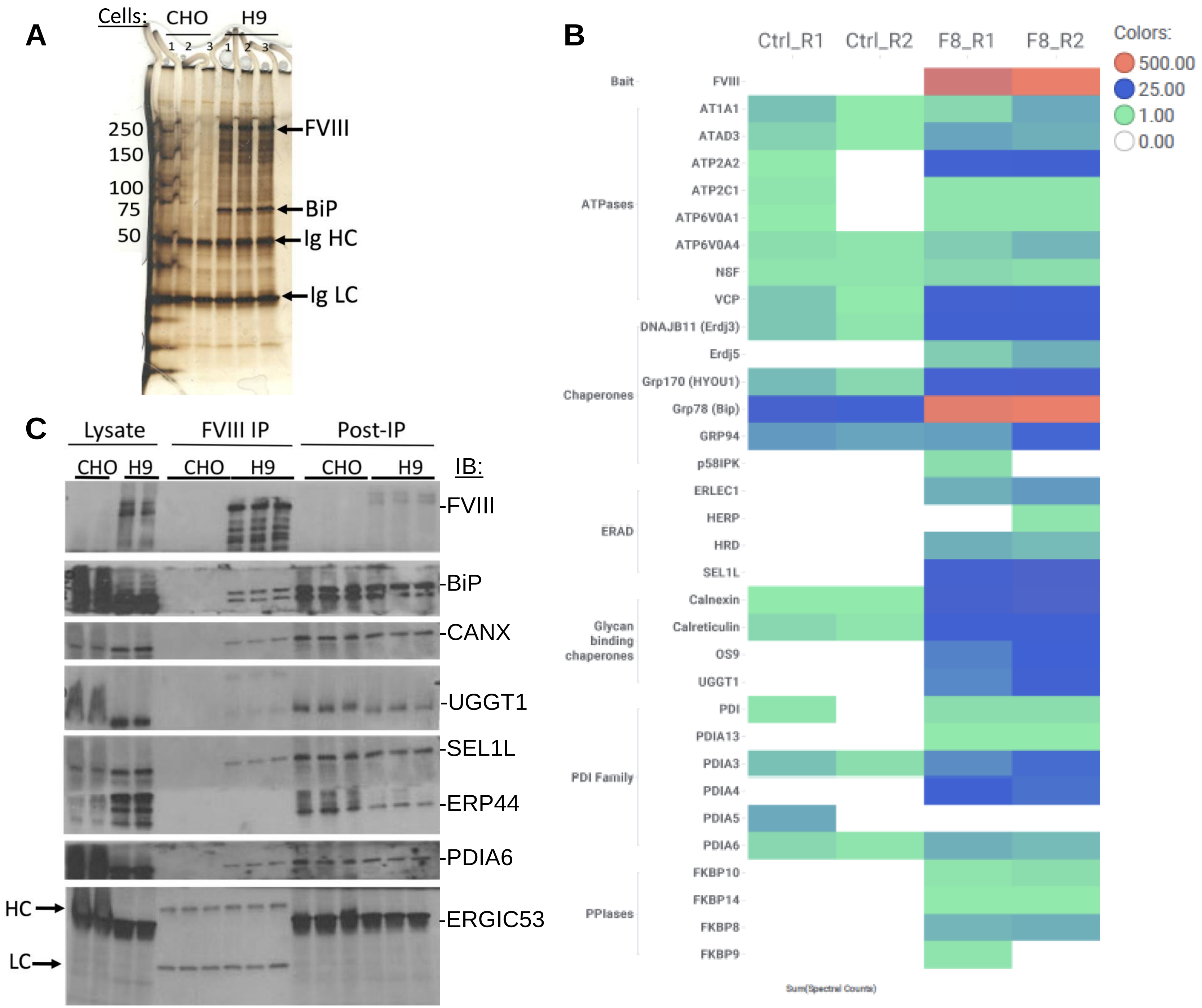
A
Cells:
CHO
H9
1 2 3 1 2 3
250
150
100
75
50
FVIII
BiP
Ig HC
Ig LC
B
Ctrl_R1
Ctrl_R2
F8_R1
F8_R2
Colors:
500.00
25.00
1.00
0.00
Bait
FVIII
ATPases
AT1A1
ATAD3
ATP2A2
ATP2C1
ATP6V0A1
ATP6V0A4
NSF
VCP
Chaperones
DNAJB11 (Erdj3)
Erdj5
Grp170 (HYOU1)
Grp78 (Bip)
GRP94
p58IPK
ERAD
ERLEC1
HERP
HRD
SEL1L
Glycan binding chaperones
Calnexin
Calreticulin
OS9
UGGT1
PDI Family
PDI
PDIA13
PDIA3
PDIA4
PDIA5
PDIA6
PPIases
FKBP10
FKBP14
FKBP8
FKBP9
Sum(Spectral Counts)
C
Lysate
FVIII IP
Post-IP
CHO H9
CHO H9
CHO H9
IB:
-FVIII
-BiP
-CANX
-UGGT1
-SEL1L
-ERP44
-PDIA6
-ERGIC53
HC
LC

Figure S7

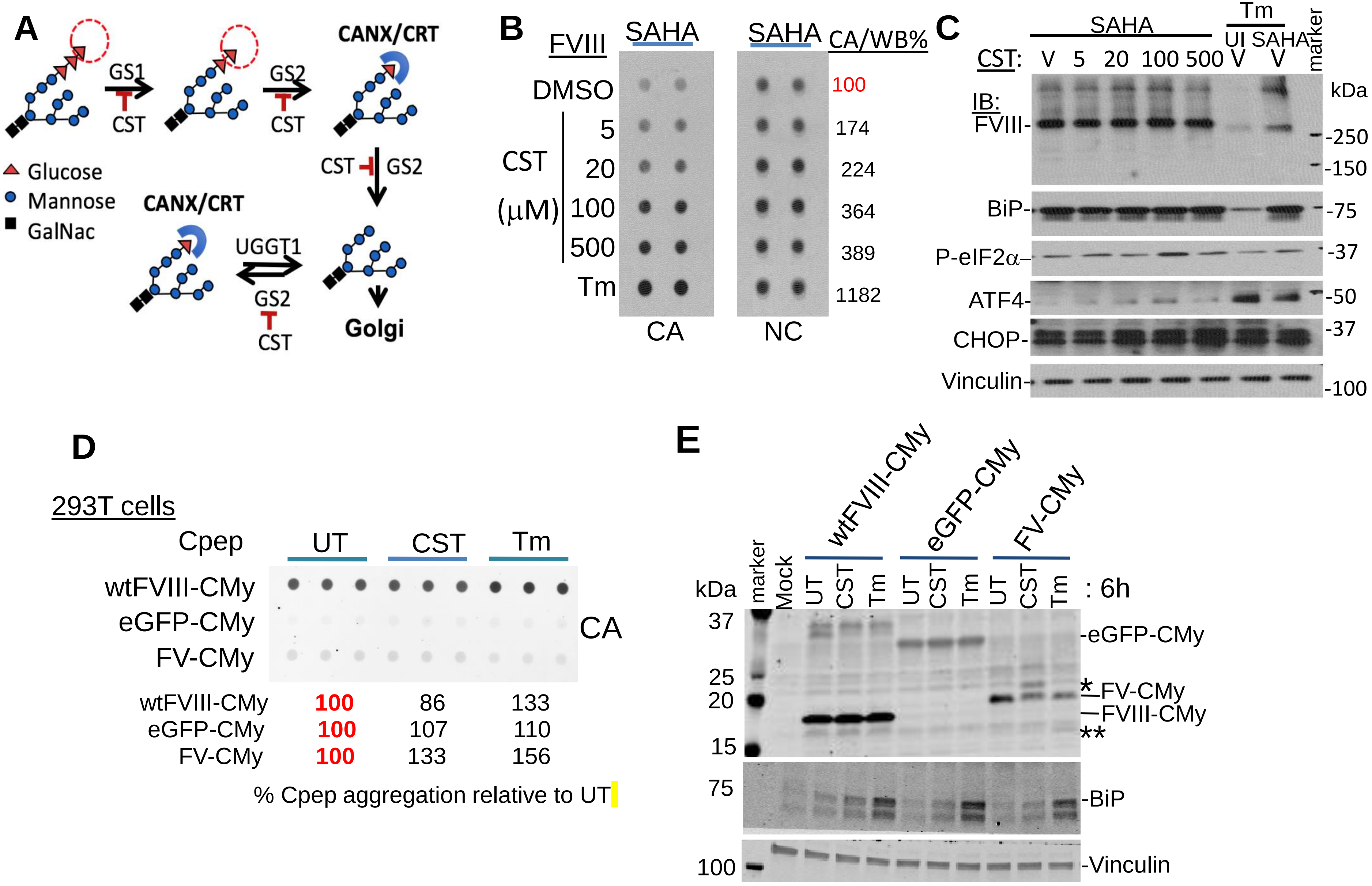


| | UT | CST | Tm |
|---|---|---|---|
| wtFVIII-CMy | 100 | 86 | 133 |
| eGFP-CMy | 100 | 107 | 110 |
| FV-CMy | 100 | 133 | 156 |

% Cpep aggregation relative to UT

### Figure S8

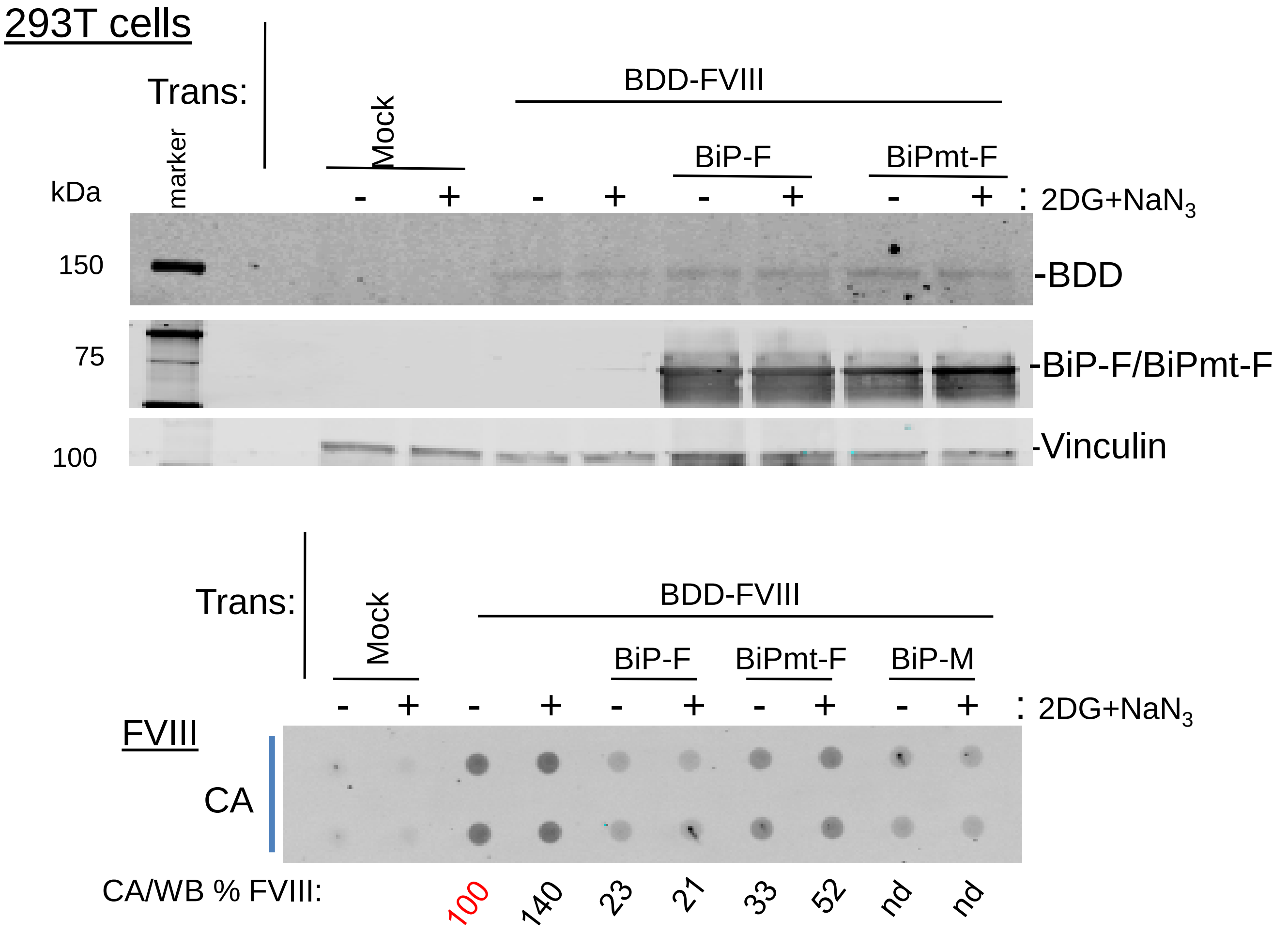
293T cells
Trans:
BDD-FVIII
Mock
BiP-F
BiPmt-F
marker
kDa
- + - + - + - + : 2DG+NaN$_3$
150
75
100
-BDD
-BiP-F/BiPmt-F
-Vinculin
Trans:
BDD-FVIII
Mock
BiP-F
BiPmt-F
BiP-M
- + - + - + - + - + : 2DG+NaN$_3$
FVIII
CA
CA/WB % FVIII:
100
140
23
21
33
52
nd
nd

Figure S9

**A**

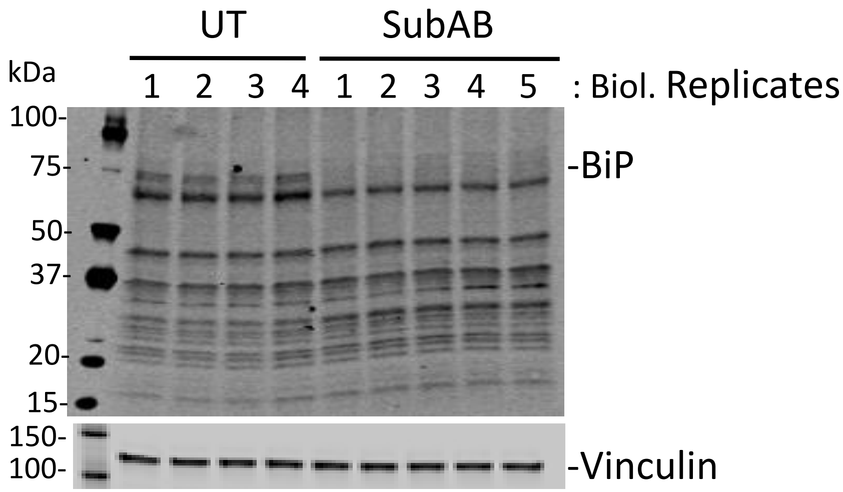
UT
SubAB
kDa
1 2 3 4 1 2 3 4 5 : Biol. Replicates
100-
75-
50-
37-
20-
15-
-BiP
150-
100-
-Vinculin

**B**

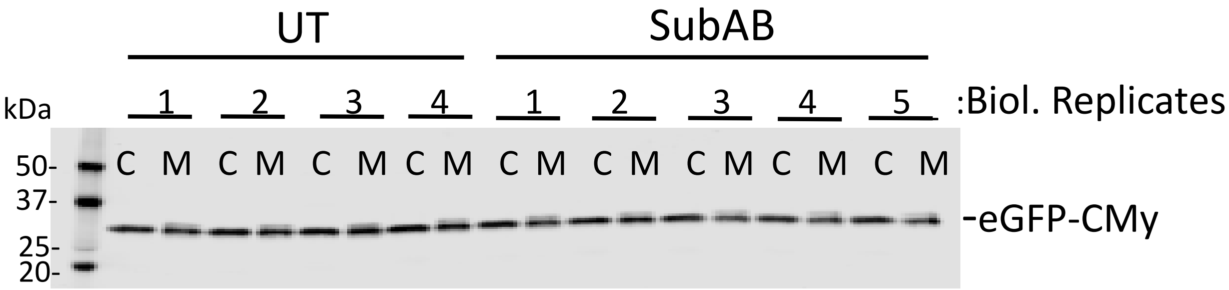
UT
SubAB
kDa
1 2 3 4 1 2 3 4 5 :Biol. Replicates
C M C M C M C M C M C M C M C M C M
50-
37-
25-
20-
-eGFP-CMy
